# The MAP3Ks DLK and LZK direct diverse responses to axon damage in zebrafish peripheral neurons

**DOI:** 10.1101/2021.07.03.450951

**Authors:** Kadidia Pemba Adula, Mathew Shorey, Vasudha Chauhan, Khaled Nassman, Shu-Fan Chen, Melissa M Rolls, Alvaro Sagasti

## Abstract

The MAP3Ks Dual Leucine Kinase (DLK) and Leucine Zipper Kinase (LZK) are essential mediators of axon damage responses, but their responses are varied, complex, and incompletely understood. To characterize their functions in axon injury, we generated zebrafish mutants of each gene, labeled motor neurons (MN) and touch-sensing neurons in live zebrafish, precisely cut their axons with a laser, and assessed the ability of mutant axons to regenerate. DLK and LZK were required redundantly and cell autonomously for axon regeneration in MNs, but not in larval Rohon-Beard (RB) or adult dorsal root ganglion (DRG) sensory neurons. Surprisingly, in *dlk lzk* double mutants, the spared branches of wounded RB axons grew excessively, suggesting that these kinases inhibit regenerative sprouting in damaged axons. Uninjured trigeminal sensory axons also grew excessively in mutants when neighboring neurons were ablated, indicating that these MAP3Ks are general inhibitors of sensory axon growth. These results demonstrate that zebrafish DLK and LZK promote diverse injury responses, depending on the neuronal cell identity and type of axonal injury.

**Significance statement:** The MAP3Ks DLK and LZK are damage sensors that promote diverse outcomes to neuronal injury, including axon regeneration. Understanding their context-specific functions is a prerequisite to considering these kinases as therapeutic targets. To investigate DLK and LZK cell-type specific functions, we created zebrafish mutants in each gene. Using mosaic cell labeling and precise laser injury we found that both proteins were required for axon regeneration in motor neurons, but, unexpectedly, were not required for axon regeneration in Rohon-Beard or dorsal root ganglion (DRG) sensory neurons, and negatively regulated sprouting in the spared axons of touch-sensing neurons. These findings emphasize that animals have evolved distinct mechanisms to regulate injury site regeneration and collateral sprouting, and identify differential roles for DLK and LZK in these processes.

## Introduction

Axon damage caused by stroke, trauma, or disease disrupts the circuits required for sensation, movement and cognition. Unlike other tissues that rely on stem cells to recover from damage, most neurons cannot be replaced, so damaged cells must themselves be repaired to restore function. Successful axon regeneration requires damage sensing, the transmission of injury signals to the nucleus, activation of pro-regenerative genes, axon guidance, and circuit reintegration (Curcio and Bradke, 2018). Our understanding of the factors regulating each of these steps is incomplete.

Mitogen-activated protein kinase kinase kinases (MAP3Ks) regulate many cellular processes, including development, differentiation, and stress responses (Craig et al., 2008). Among MAP3Ks, Dual Leucine Kinase (DLK/MAP3K12), which belongs to the Mixed Lineage Kinase (MLK) MAP3K subfamily (Gallo and Johnson, 2002), has been implicated in neuronal development (Nakata et al., 2005; Hirai et al., 2006, 2011) and axon injury responses (Hammarlund et al., 2009; Miller et al., 2009; Xiong et al., 2010; Welsbie et al., 2017, 2019) in a variety of organisms. DLK is activated by axon injury and in turn activates downstream P38 or JNK signaling cascades to induce transcription of injury response genes (Tedeschi and Bradke, 2013; Jin and Zheng, 2019). In addition to DLK itself, vertebrates have another DLK-related gene that contributes to injury responses, Leucine Zipper Kinase (LZK/MAP3K13), which has a domain analogous to the calcium-sensing domain in the worm DLK protein (Yan and Jin, 2012).

In invertebrates, activation of DLK is a major signal initiating responses to axon injury and stress. Without DLK, regenerative axon growth is eliminated in motor and sensory neurons in both *C. elegans* (Hammarlund et al., 2009; Yan et al., 2009) and *Drosophila* (Xiong et al., 2010; Stone et al., 2014). In mammals, the outcomes of DLK and LZK activation in response to axon damage are varied and context-dependent: DLK or LZK can promote neurite branching (Chen et al., 2016b), axon elongation (Shin et al., 2012), inhibition of axon regeneration (Dickson et al., 2010), axon degeneration of the distal stump (Miller et al., 2009; Summers et al., 2018), cell death (Ghosh et al., 2011; Watkins et al., 2013; Yin et al., 2017; Welsbie et al., 2019; Li et al., 2021), or microglial and astrocyte responses to injury (Chen et al., 2018; Wlaschin et al., 2018). In mice with a DLK gene-trap, downstream responses were reduced after sciatic nerve injury, and explant cultures of DRG grew shorter axons than controls (Itoh et al., 2009). Selectively eliminating DLK in neurons strongly reduced motor axon target reinnervation after sciatic nerve crush (Shin et al., 2012). Peripheral axons of DLK mutant DRG neurons, on the other hand, initiated regeneration normally, but by three days post-injury had regenerated less than controls (Shin et al., 2012).

Studying genetic regulators of axon regeneration is complicated by the complexity of in vivo experimental injuries. In nerve crush or severing models, nerve bundles contain axons of many different types of neurons, thus convoluting cell type-specific responses to injury. Moreover, axon branching and fasciculation within nerves can obscure the source of growth after incomplete injuries--without single cell resolution, it is difficult to distinguish axon growth from regenerating neurites, regenerative sprouting from unsevered branches of damaged cells, or collateral sprouting from undamaged neurons (Steward et al., 2003; Tuszynski and Steward, 2012), each of which has distinct functional implications.

To better understand the roles of *dlk* and *lzk* in axon damage responses, we created zebrafish mutants in both genes. Single-cell labeling allowed us to compare regeneration not only in different cell types, but also in central axons, peripheral axons, and specific branches of damaged axons. Our findings indicate that *dlk* and *lzk* are required redundantly for motor axon regeneration, but not for axon regeneration in RB or DRG neurons. Surprisingly, DLK inhibited sprouting of larval peripheral sensory axons, emphasizing the context-dependent multifunctionality of these axon damage sensors.

## Results

### *dlk^la231^* and *lzk^la232^* mutant zebrafish develop normal motor and sensory neurons

To study the functions of zebrafish DLK and LZK in axon regeneration, we identified the closest homologous genes to mammalian DLK and LZK in the zebrafish genome, which were located on opposite ends of chromosome 9, and generated a phylogenetic tree with the full amino acid sequences (Figure 1A). Human, mouse, and zebrafish DLK proteins share 93% sequence similarity in their kinase domain, and 95% similarity in their leucine zipper domains; human, mouse, and zebrafish LZK proteins share 97% sequence similarity in their kinase domain, 98% similarity in their leucine zipper domains, and an identical C-terminal hexapeptide involved in calcium regulation (Supplemental Figure 1).

**Figure 1.**
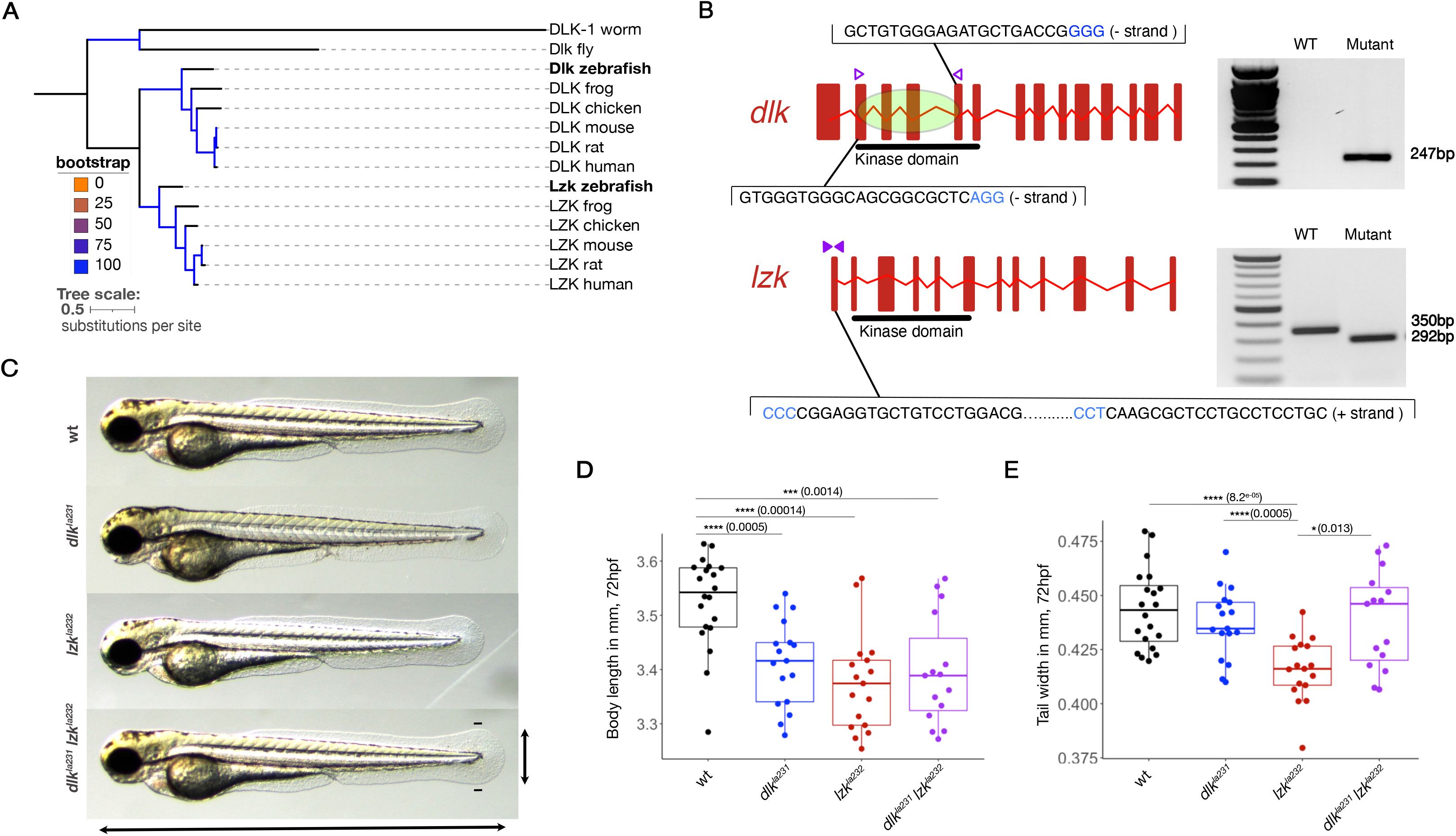
*dlk* and *lzk* zebrafish mutants. A) Phylogenetic tree of DLK and LZK orthologs. B) Genotyping of *dlk^la231^* and *lzk^la232^* CRISPR/Cas9 mutants. Left, gene structure of zebrafish *dlk* and *lzk*, with gRNA sequences. Blue indicates PAM sites. Right, DNA gel showing wt and mutant genotyping with primers indicated to the left (arrowheads). C) 48 hpf zebrafish larvae of the indicated phenotypes. D) Overlaid box and dot plots comparing animal lengths from the tip of the head to the end of the tail each genotype. E) Overlaid box and dot plots comparing tail width in each genotype. See Methods for details of statistical analyses.

We created mutations in each gene using the CRISPR/Cas9 system. Using two guide RNAs (gRNAs), we made a 3289 base pair (bp) genomic deletion in zebrafish *dlk*, which removed 519 coding base pairs, including most of the kinase domain. Using the same approach to target LZK, we created an allele with two separate deletions in exon 1 (a 28 base pair deletion and a 30 base pair deletion), placing the gene out-of-frame, upstream of the kinase domain (Figure 1B, Supplemental Figure 2). Unlike DLK mutant mice, which die perinatally (Hirai et al., 2006), *dlk^la231^*, *lzk^la232^*, and *dlk^la231^ lzk^la232^*double mutant zebrafish survived to adulthood and appeared grossly normal, similar to mutants in invertebrate DLK homologs, although both mutants were slightly smaller on average than wildtype fish at three larval stages (48 hpf, 72 hpf, and 5 dpf), indicating a developmental delay (Figure 1C-E; Supplemental Figure 3).

Zebrafish *dlk* mRNA is expressed broadly in the nervous system at early developmental stages (Thisse et al., 2004), prompting us to test if *dlk* is required for the initial development of peripheral neurons. To label motor neurons (MNs), one-cell stage embryos were co-injected with MN driver (HB9:GAL4) (Issa et al., 2011) and effector (UAS:GFP) transgenes, which drive expression of cytoplasmic GFP. Since transient transgenesis results in mosaic inheritance, we screened for animals expressing GFP in isolated MNs. To minimize morphological variability, we selected only those MNs that innervated ventral muscles (Figure 2A_B), enriching for the caudal primary (CaP) MNs (Westerfield et al., 1986). At 5 days post-fertilization (dpf) we imaged each MN and measured several morphological parameters, including axon branch tip number, and total axon length. Despite the fact that mutant animals are slightly smaller than controls, there were no significant differences in neuronal morphology between *dlk^la231^*, *lzk^la232^*, or *dlk^la231^ lzk^la232^* double mutants and wt animals (Figure 2C-D). These observations indicate that *dlk* and *lzk* are not required for the morphological development of MNs.

**Figure 2.**
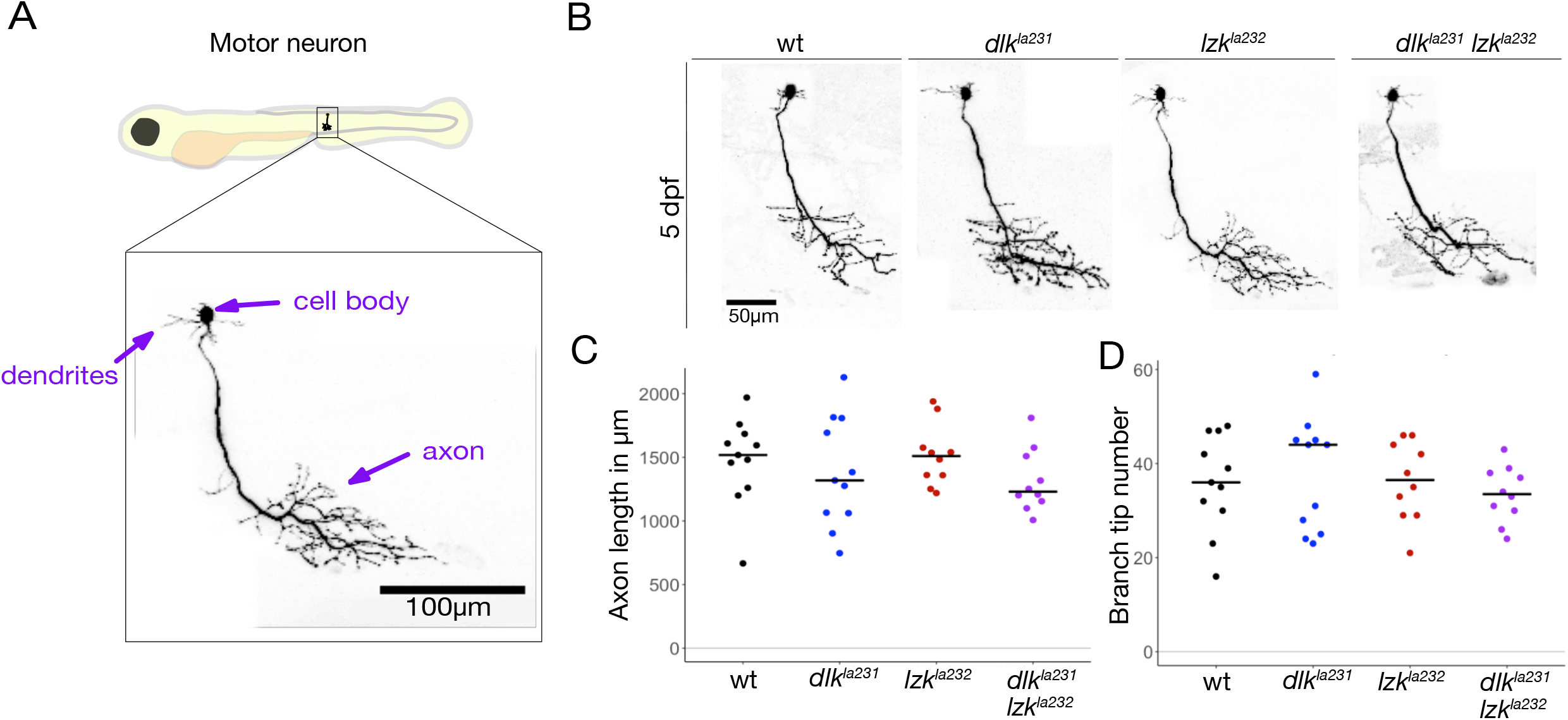
Motor neurons develop normally in *dlk^la231^* and *lzk^la232^* mutants. A) Cartoon of 5 dpf larva, showing the approximate location of the image below of a single labeled MN in a live animal. The cell body and dendrites are in the spinal cord; the axon exits the spinal cord to innervate the ventral muscles of one segment. B) Labeled MNs in each of the indicated genotypes. C) Dot plot showing lengths of MNs in each of the indicated genotypes. Bar indicates the mean. There was no significant difference between groups (because distributions were normal, groups were compared by ANOVA). D) Dot plot showing branch tip numbers of MNs in each of the indicated genotypes. Bar indicates the mean. There was no significant difference between groups. Scale bars: 100μm in A, 50μm in B.

To image larval RB touch-sensing neurons neurons (Palanca et al., 2013; Katz et al., 2021), we injected animals with a reporter driving expression of a red fluorescent protein in these cells (Isl1[SS]:Gal4;UAS:DsRed) (Sagasti et al., 2005), and screened for animals expressing this reporter in isolated RB neurons, which allowed us to unambiguously visualize the morphology of the entire neuron and distinguish central from peripheral axons. To minimize variability, we selected only tail-innervating RB neurons for analysis, since they are flat and relatively easy to trace (Figure 3A-B). As with MNs, there were no significant morphological differences between the peripheral arbors of *dlk^la231^*, *lzk^la232^*, or *dlk^la231^ lzk^la232^* double mutant and wt fish at 48 hours post-fertilization (hpf) (Figure 3C-E). At 72hpf, *dlk^la231^* and double mutant peripheral axon morphologies were also comparable to wildtype axons, but *lzk^la232^* neurons were somewhat smaller than wildtype neurons on average (Supplement Figure 4), potentially reflecting their mild developmental delay. Together these results indicate that, similar to homologs in invertebrate animals, DLK and LZK do not play major roles in initial neuronal development.

**Figure 3.**
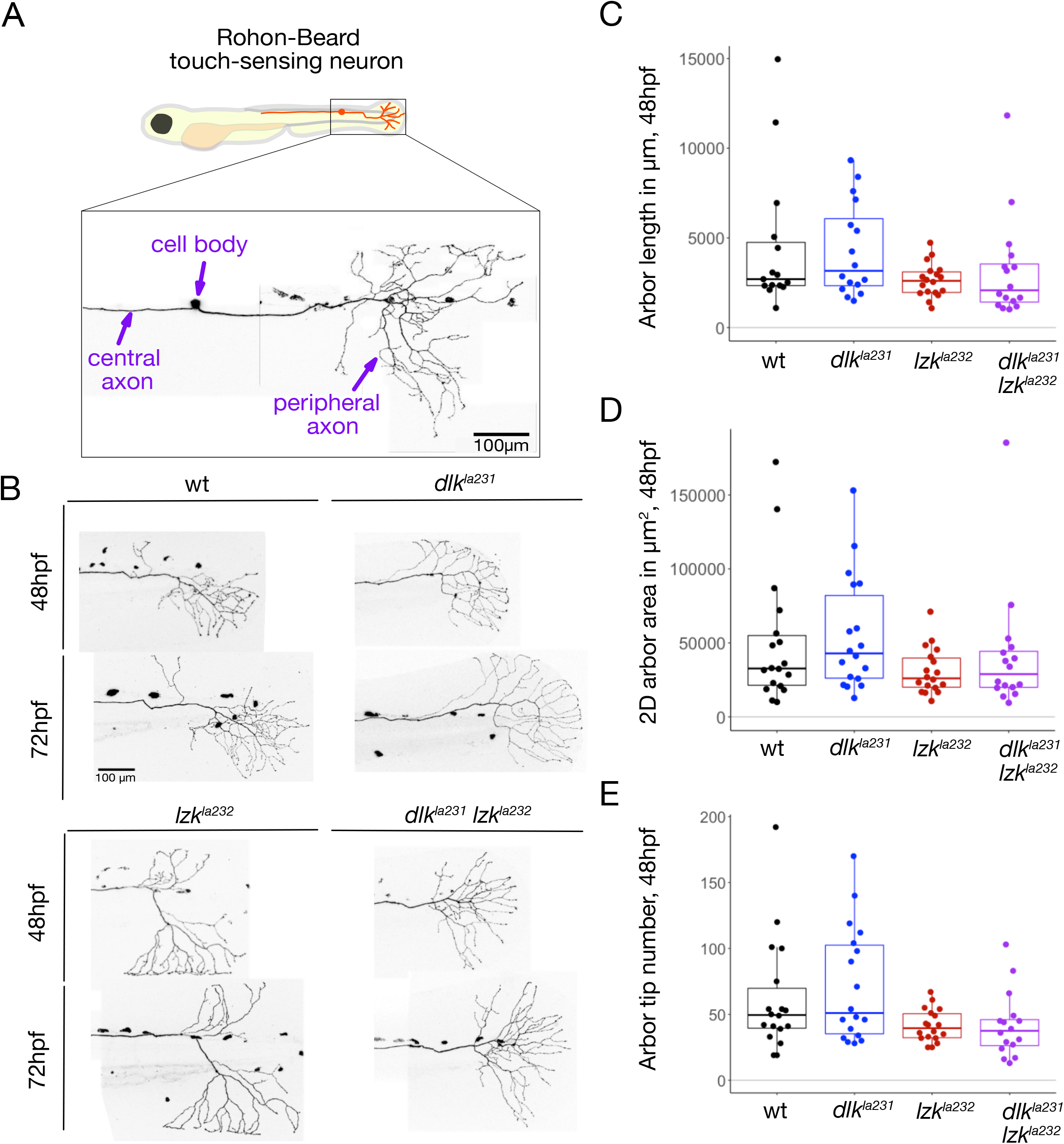
Rohon-Beard neurons develop normally in *dlk^la231^* and *lzk^la232^* mutants. A) Cartoon of 48 hpf larva, showing the approximate location of the image below of a single labeled RB neuron in a live animal. The cell body, central and peripheral axons are labeled. The cell body and central axon are in the spinal cord; the peripheral axon exits the spinal cord to arborize in the developing epidermis. B) Tail-innervating peripheral RB axon arbors of the indicated genotypes at 48 and 72 hpf. C-E) Quantification of RB peripheral axon arbor lengths (C), 2D arbor area (D), and branch tip number (E) at 48 hpf. See Methods for details of statistical analyses. Scale bars: 100μm.

### *dlk* and *lzk* are redundantly required for motor axon regeneration

Few studies have directly tested the relationship between these two closely related, potentially redundant MAP3K proteins in axon damage responses. To assess if *dlk* or *lzk* are required for motor axon regeneration in larval zebrafish, we severed axons of isolated MNs in wt, single mutant, and double mutant fish, and compared their ability to regenerate. Specifically, 5 dpf axons were severed 50 µm distal to the spinal cord exit point (Figure 4A), using a laser mounted on a 2-photon microscope (O’Brien et al., 2009b). To minimize potential contributions from extrinsic factors, only neurons in which laser axotomy caused no obvious damage or scars were used for regeneration experiments. Motor axons were assessed for regeneration 48 hours post-axotomy (hpa) (Figure 4A), as in previous studies (Rosenberg et al., 2012). Wallerian degeneration appeared to occur normally in these mutants. Motor axon regeneration was modestly reduced in *dlk^la231^* mutants, compared to wt motor axons, but strongly impaired in *dlk^la231^ lzk^la232^* double mutants (Figure 4B-C). These observations suggest that *dlk* and *lzk* are partially redundant (or genetically compensate for each other) for regeneration of larval motor axons.

**Figure 4.**
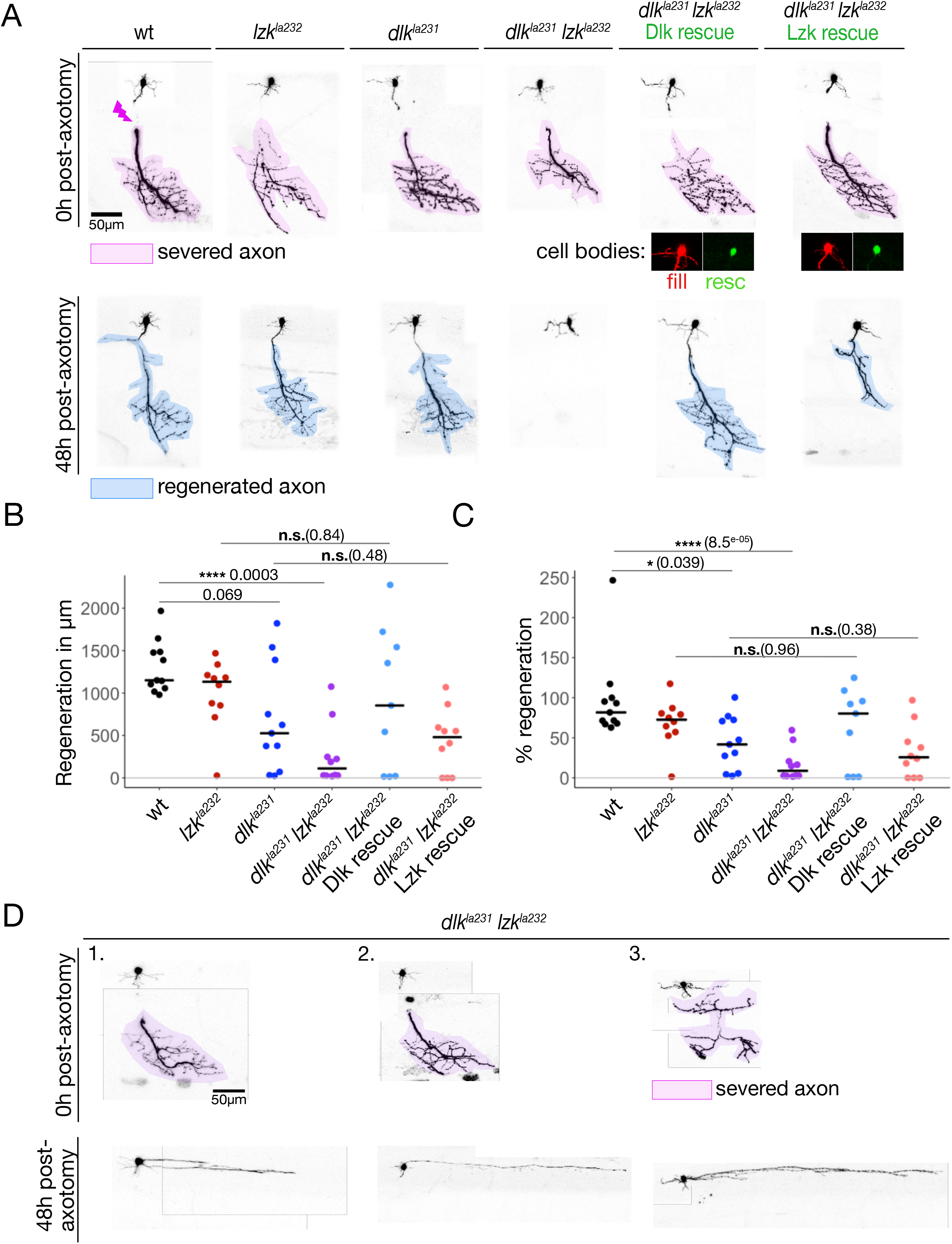
Motor neuron regeneration is impaired in *dlk^la231^ lzk^la232^* mutants. A) Top: 5 dpf motor axons immediately after axotomy in the indicated genotypes. Lightning bolt indicates axotomy site. Magenta highlights the separated distal stump that will degenerate. Rightmost panels show neurons expressing rescue cDNAs; expression of rescue transgenes in cell bodies is shown below. Bottom: Same neurons 48 hours post-axotomy. Blue highlights regenerated axons. B-C) Dot plots showing total regenerated length in each genotype (B) and the percentage of the original axon length regenerated (C). Bar indicates the mean. D) Dendrite overgrowth phenotype following failure of axon regeneration in *dlk^la231^ lzk^la232^* mutants. See Methods for details of statistical analyses.

Intriguingly, we observed that among *dlk^la231^ lzk^la232^* double mutant MNs displaying severe regeneration deficits, ∼28% (8 out of 29) grew extremely long neurites within the spinal cord, which are likely dendrites (Figure 4D), a phenotype not seen in individual *dlk^la231^* or *lzk^la232^* mutants. This observation indicates that *dlk^la231^ lzk^la232^* double mutant axons do not lack growth potential, but are specifically impaired in axon regeneration.

To determine if *dlk* or *lzk* are required cell-autonomously for motor axon regeneration, we attempted to rescue their regeneration defects by expressing *dlk* and *lzk* cDNAs specifically in these neurons. Since *dlk* and *lzk* were required redundantly for motor axon regeneration, we expressed each cDNA separately in *dlk^la231^ lzk^la232^* double mutants. Strong overexpression of these cDNAs with the Gal4/UAS system was toxic to neurons, causing dysmorphic axons, spontaneous axon degeneration, and cell death (not shown). We therefore expressed lower levels by creating bicistronic transgenes directly under a MN-specific promoter (HB9:DLK*-*T2A-GFP and HB9:LZK-T2A-GFP), which were co-injected with transgenes that strongly drive RFP expression throughout the cytoplasm (HB9:Gal4 and UAS:DsRed). Only cells that clearly expressed the rescue transgene (i.e., GFP+ cells), and had overtly normal axons, were used for these experiments (Figure 4A). Expressing each cDNA improved axon regeneration to levels comparable to the corresponding single mutant (e.g., *dlk^la231^ lzk^la232^* neurons with *dlk* cDNA were comparable to *lzk^la232^* mutants, Figure 4B-C). However, the difference in regeneration between rescued neurons and double mutants did not reach significance, likely due to variability in the assay. Nonetheless, these results are most consistent with DLK and LZK proteins acting cell-autonomously to promote motor axon regeneration, similar to DLK’s well documented role in axon regeneration in worms, flies, and mice.

### *dlk* and *lzk* are not required for RB central or peripheral axon regeneration

RB neurons are bipolar or pseudo-unipolar, with a receptive peripheral axon that innervates the epidermis and a central axon that connects to downstream circuits in the spinal cord (Palanca et al., 2013; Katz et al., 2021). Unlike in mammals, fish can often regenerate axons in the central nervous system (Rasmussen and Sagasti, 2016). To visualize RB neurons, we used the *Islet1*(ss) enhancer, which drives expression in all touch-sensing neurons (Higashijima et al., 2000; Sagasti et al., 2005). To identify a time point for regeneration experiments, we characterized the structure of tail-innervating RB neuron arbors as development progressed, with the aim of finding a stage when RB neurons were morphologically stable, but still expressed the transgene strongly. These analyses led us to choose 48hpf for axotomies, since approximately ⅔ of RB peripheral arbors in the tail had attained a stable branching pattern (i.e., new branches were no longer added) at this time point, although they continued to grow bigger via scaling growth.

To test if *dlk* or *lzk* are required for the regeneration of zebrafish RB central axons, we transiently labeled isolated RB neurons in the tail, laser severed their ascending central axons 200 μm from the cell body, and measured the total length of regenerated axons at 24 hpa (Figure 5A-B). We were careful to use the minimal effective laser power for axotomy, since we observed that even moderate damage and scarring in the spinal cord impedes regeneration. Surprisingly, RB central axons in *dlk^la231^*, *lzk^la232^*, and *dlk^la231^ lzk^la232^* double mutants regenerated similar to wt neurons (Figure 5B). In fact, some central axons regenerated more on average than wt axons, although this effect did not reach significance. These data indicate that, unlike zebrafish motor axons, or central axons of sensory neurons in invertebrates and mice, central axons of RB neurons in zebrafish larvae do not require *dlk* or *lzk* for regeneration.

**FIgure 5.**
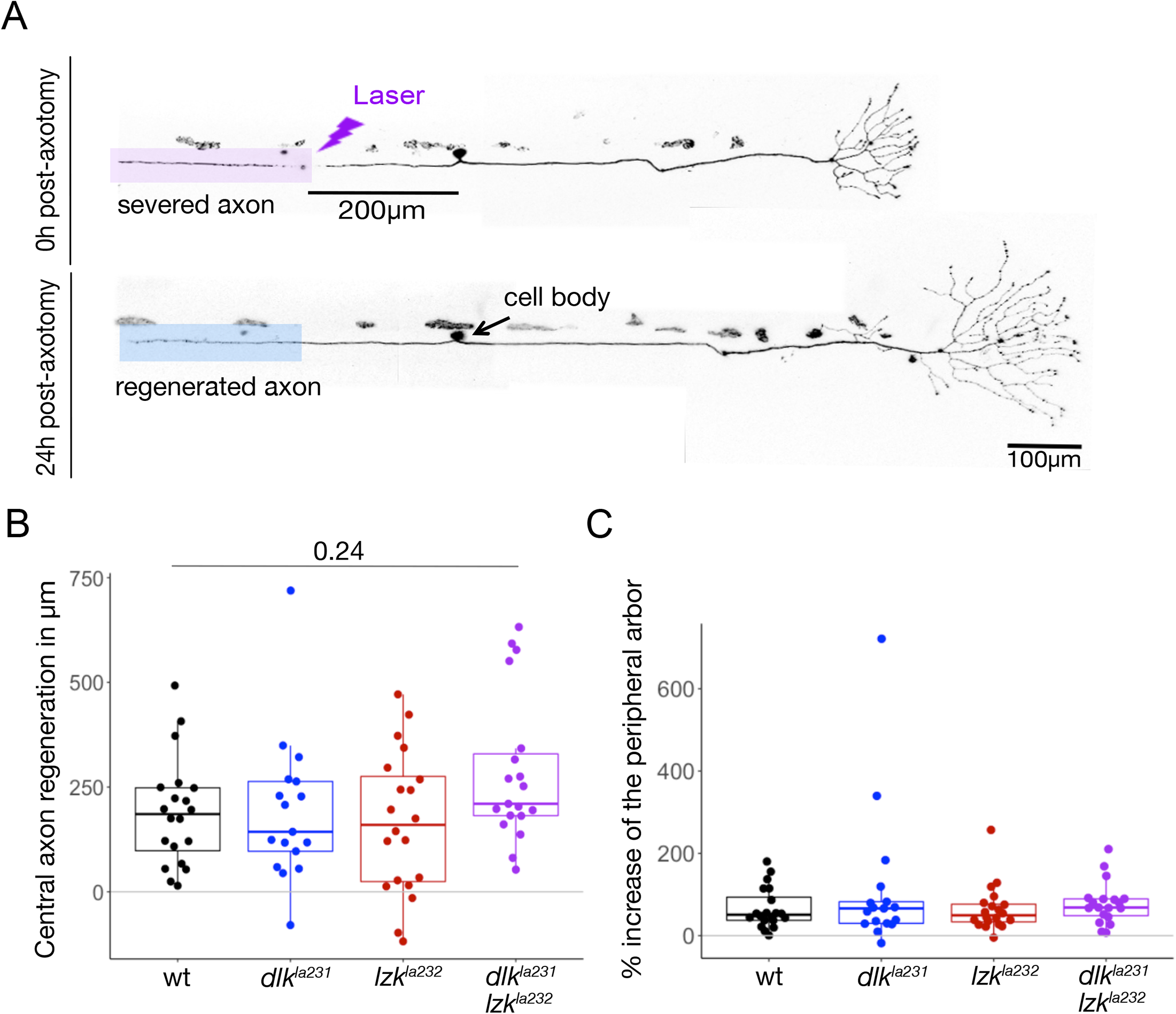
RB central axons regenerate in *dlk^la231^ lzk^la232^* mutants. A) Top: 48 hpf RB axon immediately after axotomy. Magenta highlights the separated distal stump that will degenerate. Bottom: Same neuron 24 hours post-axotomy. Blue highlights the regenerated axon. B) Overlaid box and dot plots showing central axon length regenerated in the indicated genotypes. C) Overlaid box and dot plots showing growth of peripheral arbors following central axotomy in the indicated genotypes. See Methods for details of statistical analyses. Scale bar: 100μm.

To test if *dlk* or *lzk* are required for the regeneration of RB peripheral axons, we labeled isolated tail-innervating RB neurons, removed the entire peripheral arbor by severing it at the first peripheral branch point at 48hpf, and measured regeneration at 24 hpa (Figure 6A). (When a neuron had two arbors separately innervating the skin, both were removed). Although a few axons in each genotype failed to regrow, most severed peripheral axons in single and double mutants regenerated comparable to wt, whether measured as total axon growth or as percent regeneration of the severed arbor (Figure 6B-C). Although all genotypes regenerated on average similar size arbors (Figure 6B), there were some differences in regeneration between different mutant genotypes when measured as percent regeneration of the original arbor (Figure 6C), reflecting that each group had different sized arbors to begin with. Thus, neither central nor peripheral RB axons require DLK or LZK for regeneration.

**Figure 6.**
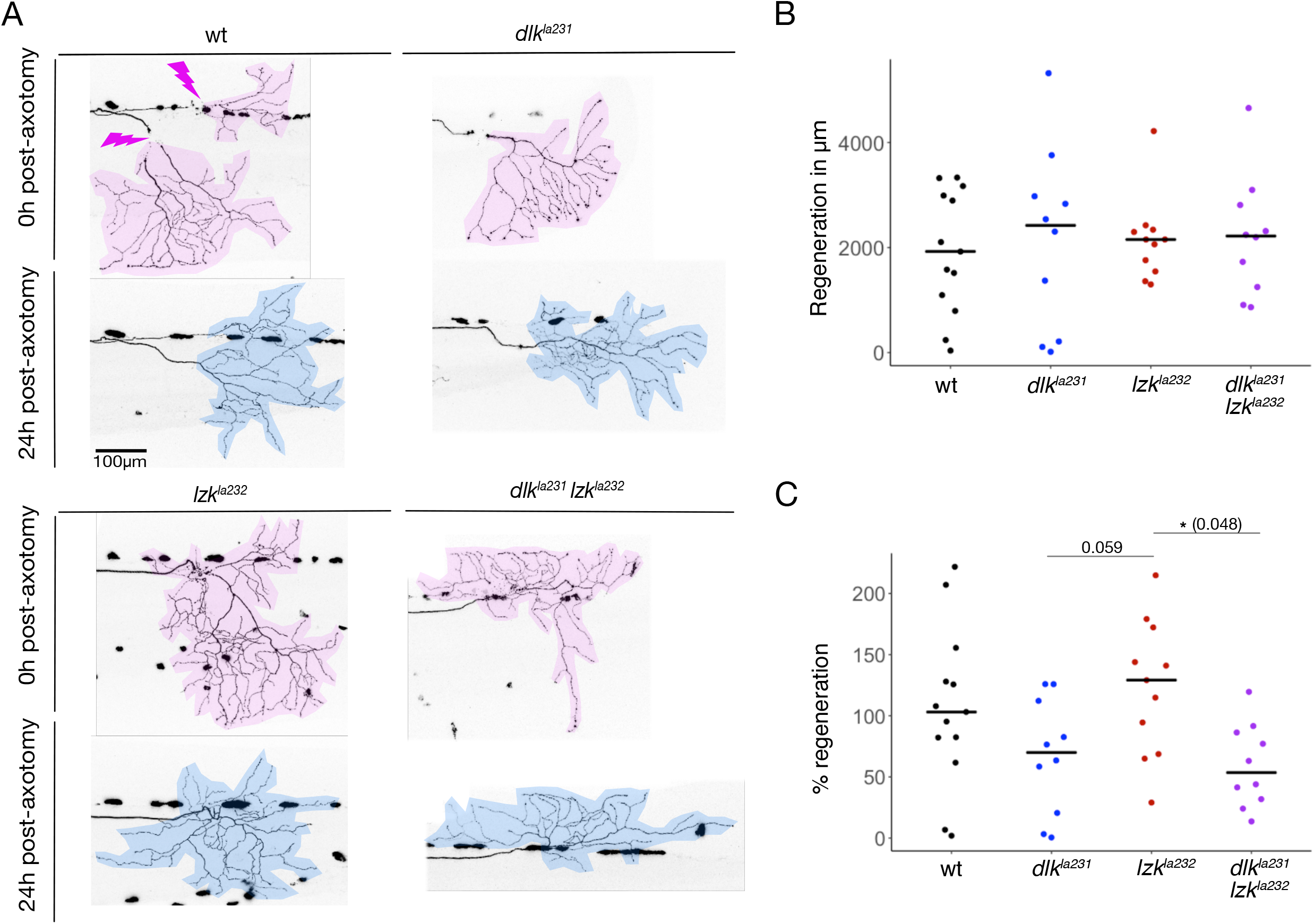
RB peripheral axon arbors regenerate in *dlk^la231^ lzk^la232^* mutants. A) Top panels: 48 hpf RB peripheral axon immediately after axotomy in the indicated genotypes. Magenta highlights the separated distal stump that will degenerate. Bottom panels: Same neurons 24 hours post-axotomy. Blue highlights the regenerated axon. B-C) Dot plots showing total regenerated length in each genotype (B) and the percentage of the original axon length regenerated (C). Bar indicates the mean. See Methods for details of statistical analyses. Scale bar: 100μm.

### *dlk* and *lzk* are not required for DRG neuron peripheral axon regeneration in adult zebrafish

The finding that DLK and LZK are not required for regeneration of RB axons was surprising, since MNs in the same genetic background failed to regenerate and DLK signaling has been implicated in regeneration of another sensory neuron type, DRG neurons in culture and adult mice (Itoh et al., 2009; Shin et al., 2012). We therefore considered the possibility that RB neurons, which are replaced by DRG neurons over the course of development (Reyes et al., 2004; Rasmussen et al., 2018), might use a repair strategy different from neurons that persist into adulthood. To determine if DRG neurons require DLK and LZK for regeneration, we imaged them in a line stably expressing mCherry in sensory neurons (P2rx3a:LexA,4xLexAop:mCherry^la207^), in wildtype, *dlk^la231^, lzk^la232^*, and *dlk^la231^ lzk^la232^* double mutant animals. Some genotypes were crossed to the *casper* mutant background, which lacks pigmentation (White et al., 2008) to facilitate imaging (see Methods for details of transgenic genotypes). We first examined regeneration of adult (8-11 month old) DRG neurons by severing sensory nerves immediately above and under a scale (Figure 7), as these nerves were easily accessible for injury with a pulsed UV laser (Rasmussen et al., 2018). Adult fish were intubated during injury and subsequent imaging sessions (Shorey et al., 2021). By 24 hours after axotomy, neurites distal to the cut site had degenerated, and by 96 hpa, scales were again fully innervated, comparable to pre-axotomy conditions, in wt, *dlk^la231^, lzk^la232^*, and *dlk^la231^ lzk^la232^* double mutants (n= 10 wt; n= 7 *dlk^la231^*; n= 5 *lzk^la232^*; n= 5 double mutants) (Figure 7).

**Figure 7.**
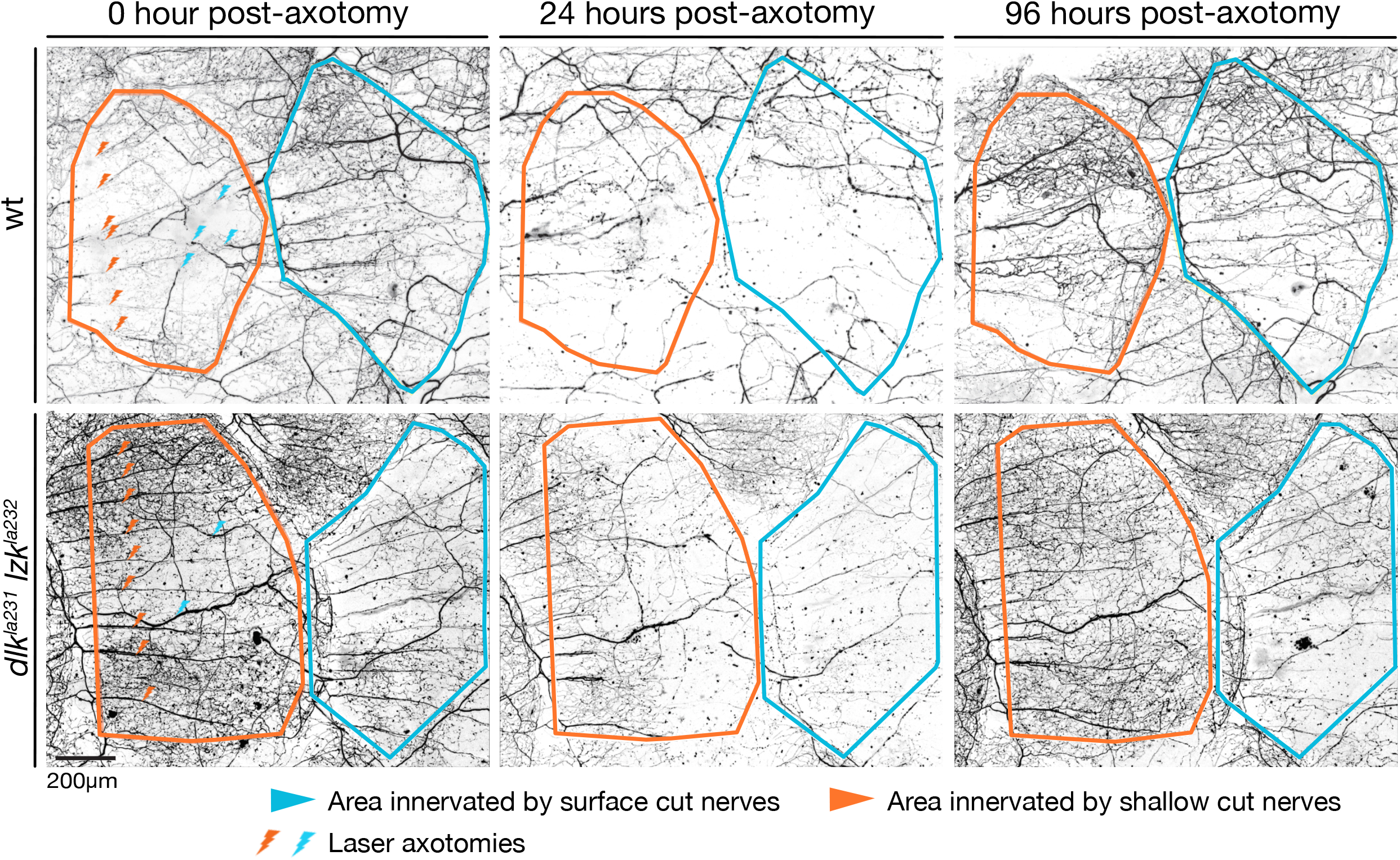
DRG axons innervating the adult scale regenerate in *dlk^la231^ lzk^la232^* mutants. Images of DRG neurons innervating the epidermis above scales in adult (8-11 month old) zebrafish in indicated genotypes at 0 hours post-axotomy (top shows immediately after axotomy, bottom shows immediately before axotomy). Lightning bolts indicate axotomy sites of individual nerves growing into scales. Nerves innervating anterior scales (orange) are above the scale, whereas nerves innervating posterior scale (blue) are below the anterior scale. Legend refers to the image on the right. Most axons innervating these scales have degenerated by 24hpa (middle column), and reinnervated scales by 96hpa (right columns). Scales were re-innervated in all wt (n=10), *dlk^la231^*(n=7), *lzk^la232^*(n=5), and *dlk^la231^ lzk^la232^* mutants (n=5). Scale bar: 200μm.

A previous study showing reduced regeneration of DRG peripheral axons in DLK mutant neurons monitored regrowth after injury closer to the cell body (Shin et al., 2012), so we considered the possibility that surface damage of sensory axons might be so common that it does not require DLK/LZK signaling. We thus severed DRG nerves close to the ganglion, using 2-photon laser surgery in 4-5 week old juvenile fish (Figure 8A). At this age, DRG sensory arbors on the scale surface have not yet achieved a fully mature morphology, but tail-innervating neurites were indistinguishable from those of adults. We specifically cut the peripheral nerve exiting the most caudal tail-innervating DRG (potentially the equivalent of the sacral ganglion nerve) at approximately 100 microns from the ganglion, resulting in the complete loss of tail fin innervation along the dorsal-most fin rays (Figure 8B). Surprisingly, the nerve regenerated robustly in both wt and *dlk^la231^ lzk^la232^* double mutants (n= 5 wt; n= 6 double mutants); by 96hpa regenerated axons had reached the end of the tail (Figure 8B). Thus, unlike motor axons, sensory axons of RB and DRG neurons do not require LZK and DLK for regeneration.

**Figure 8.**
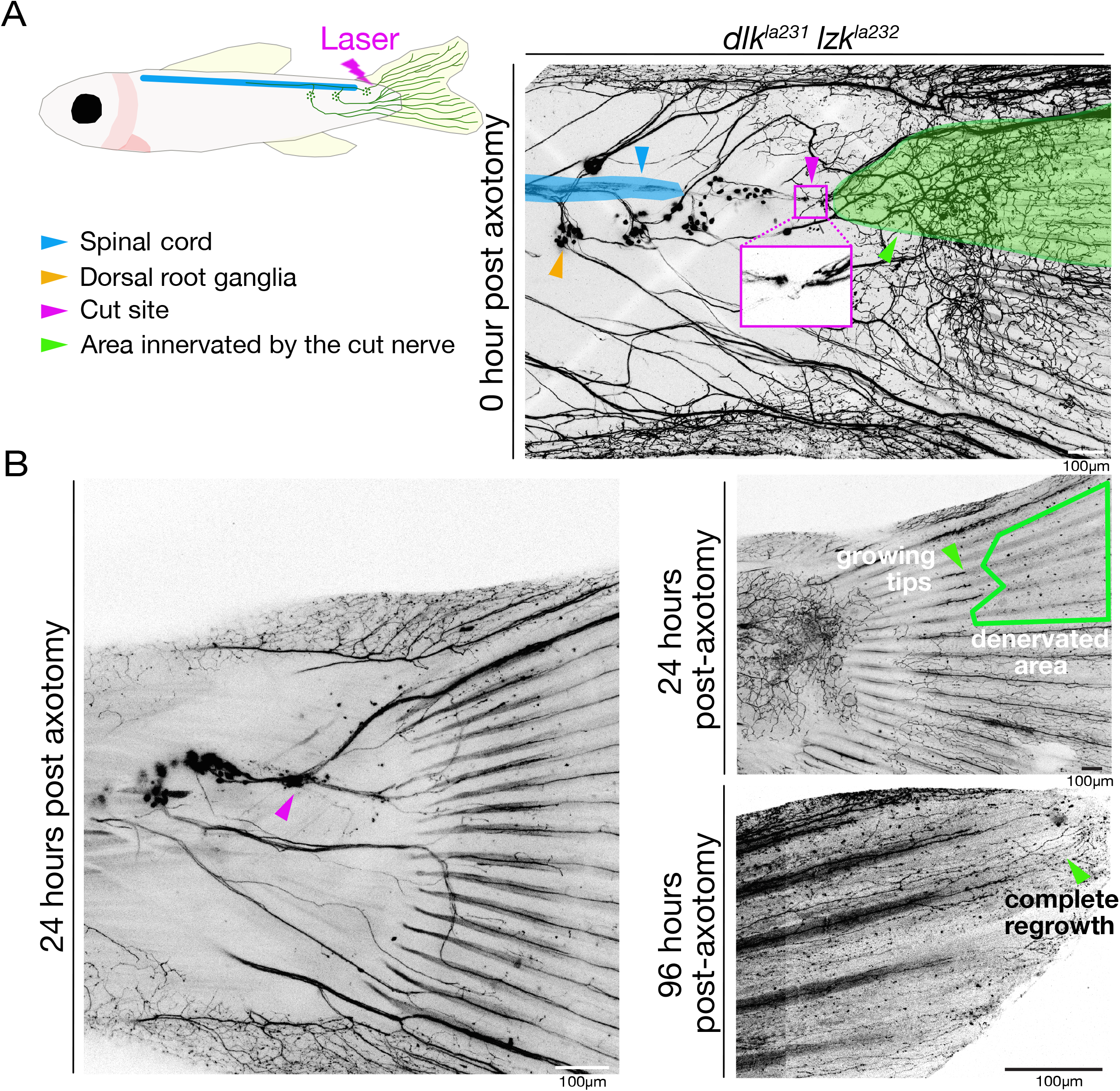
DRG axons innervating the juvenile tail regenerate in wildtype and *dlk^la231^ lzk^la232^* mutants. A) Top left: Cartoon of juvenile casper fish, showing tail-innervating DRGs in P2rx3a:LexA;4xLexAop:mCherry^la207^ transgenic fish. Lightning bolt indicates axotomy site. Blue indicates spinal cord. Legend refers to the image on the right. Top Right: 2-photon overview showing DRG cell body position, sensory nerve layout and an example of axotomy location in a 4-5 week-old zebrafish. Inset shows the axotomy site of the caudal-most DRG peripheral nerve. Green highlights separated arbors of the severed DRG nerve, which will degenerate after axotomy. B) Bottom left: Homozygous *dlk^la231^ lzk^la232^* mutant experimental animal 24 hours post-axotomy showing that axons have grown past the axotomy site. Bottom right: Fin of the same animal 24 and 96 hours post-axotomy at different magnifications. At 24 hpa, axons have grown into the fin, but have not yet reached the fin tip. By 96 hpa, axons have reached the fin tip. Axons regenerated in all wt (n=5) and all *dlk^la231^ lzk^la232^* mutants (n=6). Scale bars: 100μm.

### Spared branches of damaged RB peripheral axons sprout excessively in *dlk^la231^ lzk^la232^* double mutants

Axon regeneration from severed neurites, regenerative sprouting from spared branches of damaged axons, and collateral sprouting from undamaged axons are distinct processes (Steward et al., 2003; Tuszynski and Steward, 2012). Our single RB neuron labeling technique provided an opportunity to ask if DLK or LZK play distinct roles in these processes. To compare the behavior of severed and spared branches of a damaged axon, we severed RB peripheral axons at the second branch point, sparing the first branch (Figure 9A-B). Surprisingly, partially axotomized peripheral axons in *dlk^la231^* and *lzk^la232^* mutants regenerated more than wt axons--in other words, mutant axons gained more total peripheral axon length than wt axons after axotomy (Figure 9C). By separately measuring growth in the spared and axotomized branches, we found that regenerative sprouting contributed virtually all of the excess growth, rather than growth from the injury site (Figure 9D-E, Supplemental Figure 5). The *dlk^la231^* mutant phenotype was stronger than the *lzk^la232^* mutant phenotype, but excess growth was more pronounced in *dlk^la231^ lzk^la232^* double mutants than in either single mutant. These results indicate that *dlk* and *lzk* inhibit peripheral axon growth specifically in spared branches of damaged axons. To test if this effect is compartment-specific, we measured the growth of peripheral axons after severing the central axon of the same neuron. Cutting central axons did not promote growth of spared peripheral axons preferentially in mutants, indicating that growth inhibition by DLK and LZK occurs locally, within a compartment (Figure 5C).

**Figure 9.**
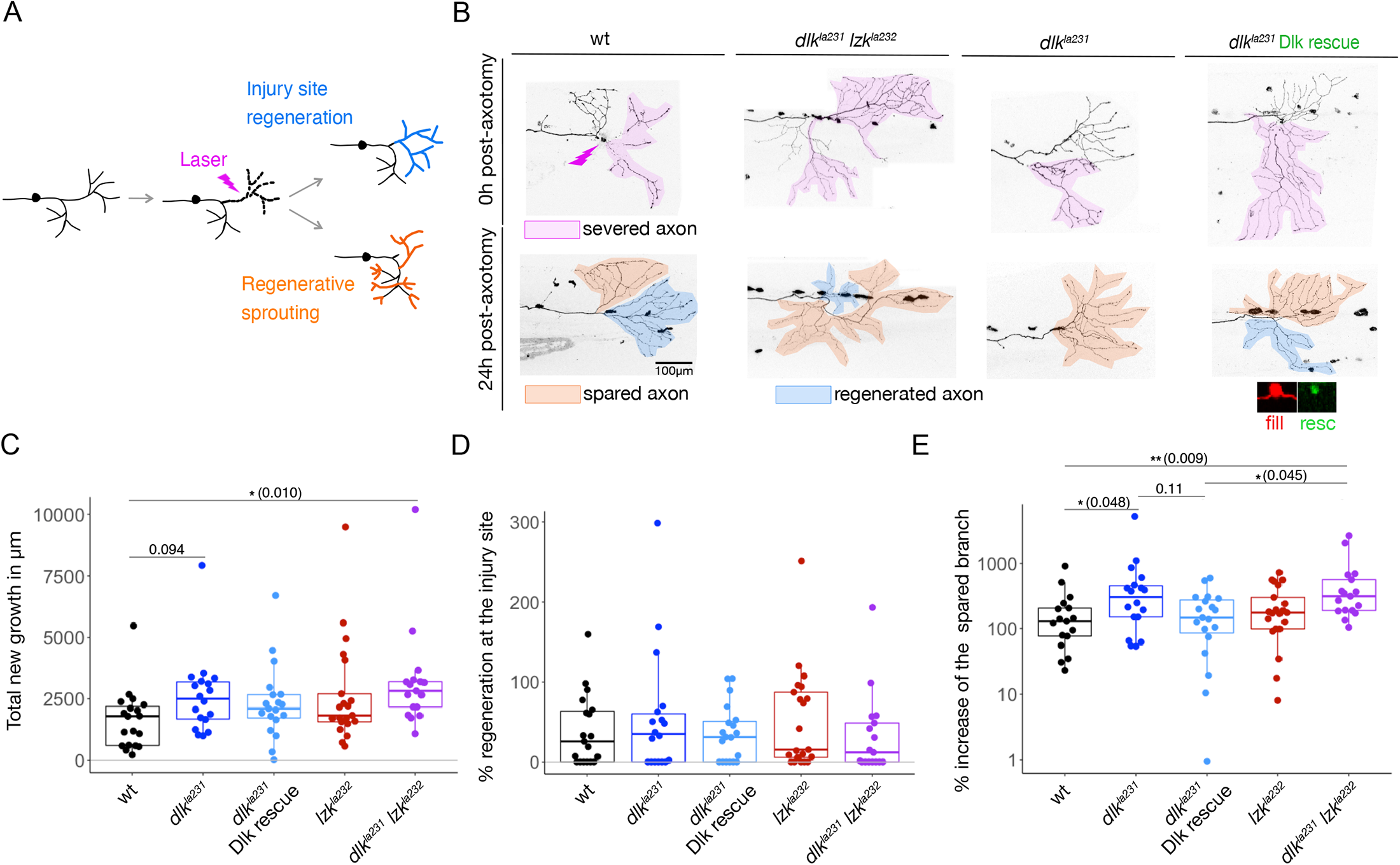
Spared arbors of damaged RB neurons sprout excessively in *dlk^la231^ lzk^la232^* mutants. A) Cartoon of partial RB peripheral axotomy assay, which differentiates between regeneration from the cut site and regenerative sprouting from spared branches. B) Top: 48 hpf RB axons immediately after axotomy in the indicated genotypes. Lightning bolt indicates axotomy site. Magenta highlights the separated distal stump that will degenerate. Rightmost panel shows a neuron expressing rescue cDNA; expression of the rescue transgene in the cell body is shown below. Bottom: Same neurons 24 hours post-axotomy. Blue highlights regenerated axons; orange highlights the spared branch. C-E) Box and dot plots showing total new growth, including both from the axotomy site and spared branch (C), percent regeneration from just the injury site (D), and percent increase of the spared branch (E). See Methods for details of statistical analyses. Scale bar: 100μm.

To test if DLK acts cell-autonomously to inhibit regenerative sprouting, we expressed *dlk* cDNA in *dlk^la231^* mutants. Like with MNs, strong overexpression of *dlk* cDNA with the Gal4/UAS system was toxic to RB neurons, causing cell death by 24hpf (not shown). As a result, we expressed lower levels of the cDNA (Crest3:DLK*-*T2A-GFP, co-injected with Isl1[SS]:Gal4 and UAS:DsRed) in RB neurons. Expression of the *dlk* cDNA modestly reduced axon sprouting, when measured as percent of the initial arbor size regenerated. This effect did not reach significance when compared to the *dlk^la231^* mutants (Figure 9C-E), but the rescued single mutants were significantly different from *dlk^la231^ lzk^la232^* double mutants. However, when measured as total axon length regenerated there was no apparent rescue with *dlk* cDNA. These ambiguous results may reflect the difficulty of achieving the appropriate level of cDNA expression for rescue, but could also suggest that these proteins do not act strictly cell-autonomously to limit regenerative sprouting of spared branches in RB neurons.

### *dlk* and *lzk* are general inhibitors of collateral sprouting in RB neurons

Given our unexpected finding that spared branches of damaged RB neurons grew more in mutants than wt animals, we wondered if increased axon growth in *dlk^la231^* and *lzk^la232^* mutants was specific to injured axons, or if sensory axon growth was generally disinhibited in these mutants. Since axon growth during development is limited by repulsive tiling interactions (Sagasti et al., 2005; Grueber and Sagasti, 2010), increased growth potential in *dlk^la231^ lzk^la232^* mutants could be masked by tiling. We therefore compared peripheral axon growth in mutants and wt animals in which we relieved axon tiling by ablating an entire trigeminal ganglion, since larval trigeminal sensory neurons are similar to larval RB neurons (Gau et al., 2013; Palanca et al., 2013). Ablating a trigeminal ganglion early in development (24hpf) allows axons to grow into the denervated side of the head significantly more than in non-ablated control animals, since axon arbors are not limited by their contralateral counterparts (Sagasti et al., 2005). However, this growth ability diminishes by 78hpf (O’Brien et al., 2009a). To reduce variegation of the reporter (i.e., silencing in some cells), we selected homozygous transgenic embryos (Isl1[SS]:Gal4;UAS:DsRed) in which most, if not all, trigeminal cell bodies were labeled. Although axons from each ganglion do not stop abruptly at a sharp midline, they rarely reach the contralateral eye in wildtype animals. 24 hours after ganglion ablation, axons of the spared ganglion grew markedly further into the denervated side of the head in *dlk^la231^ lzk^la232^* than in wt animals (Figure 10A). Measuring the total axon length that grew over the contralateral eye revealed that *dlk^la231^* and *lzk^la232^* inhibit collateral sprouting of RB touch-sensing neurons, even in uninjured neurons (Figure 10B) .

**Figure 10.**
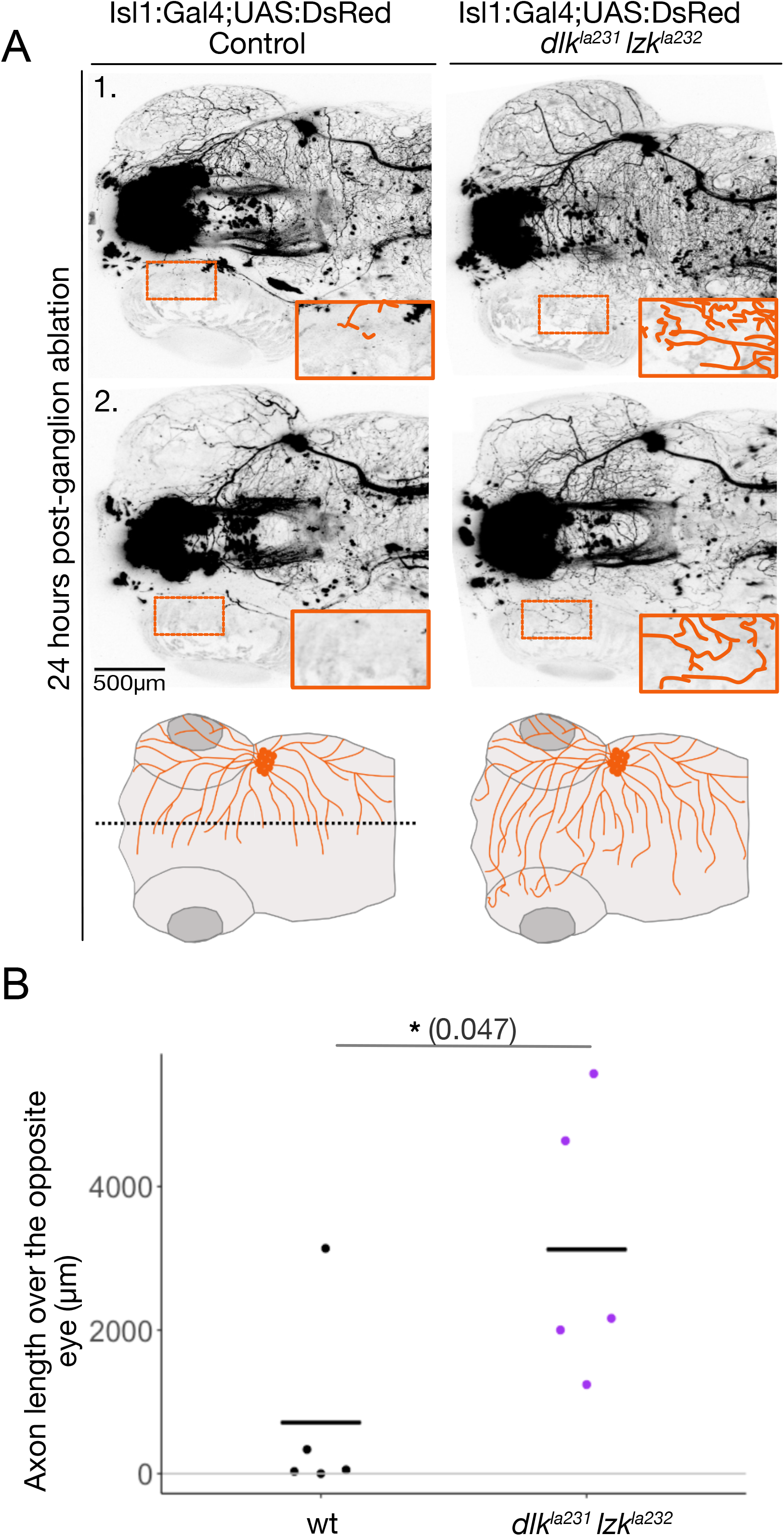
Trigeminal axons grow excessively in *dlk^la231^ lzk^la232^* mutants after ablation of the contralateral ganglion. A) Images and cartoon of trigeminal axons in zebrafish heads at 78 hpf. Confocal images show two separate examples of wt (left) and *dlk^la231^ lzk^la232^* mutant (right) fish. Insets magnify a region over the eye, with axons traced in orange. Bottom shows cartoon depictions of the result. B) Dot plot showing total axon length that grew over the contralateral eye. Bar indicates the mean. Scale bar: 500μm.

## Discussion

This study establishes and characterizes zebrafish mutants in the *dlk* and *lzk* genes, complementing existing worm, fly, and mouse models to study the function of these critical neuronal injury regulators (Jin and Zheng, 2019). Zebrafish offers powerful advantages over other vertebrate models, including the ability to label single neurons of different types and precisely injure them with laser axotomy, thus making it possible to distinguish the responses of different neurite branches to injury. Using this approach, we found that *dlk* and *lzk* were required cell-autonomously, and partially redundantly, for MN regeneration in larval zebrafish. By contrast, *dlk* and *lzk* were not required for axon regeneration in larval RB touch-sensing neurons or adult DRG neurons. However, these kinases negatively regulated the sprouting of spared sensory neuron peripheral arbors, both within the injured neuron and in uninjured neighboring neurons. These findings reveal cell-type specific *dlk* and *lzk* functions and highlight the mechanistic differences between different kinds of regenerative growth, which can be promoted or inhibited by the same signaling molecules.

Invertebrates only have one DLK-related gene, but the existence of DLK’s close relative LZK in vertebrates raises the possibility that these MAP3Ks act redundantly (Jin and Zheng, 2019). Despite their similarity, however, DLK lacks a key calcium-binding domain found in LZK and in invertebrate DLK homologs, suggesting that these kinases may be activated in different ways. Studies using optic nerve crush and traumatic brain injury (TBI) models found that inhibition of both *dlk* and *lzk* offered the strongest protection from cell death (Welsbie et al., 2019), indicating that they play overlapping roles in promoting injury-induced death. By contrast, other studies suggest distinct, cell-type-specific gene functions, including LZK’s roles in activating astrocytes (Chen et al., 2018). We directly addressed the potential for redundancy between DLK and LZK in zebrafish axon regeneration by comparing each mutant to double mutants. In motor axons, *dlk^la231^* mutants had a modest regeneration defect, but *dlk^la231^ lzk^la232^* double mutants had a strong defect, suggesting partial redundancy. *dlk^la231^* has an in-frame deletion of the kinase domain, so it is unlikely to trigger genetic compensation (Rossi et al., 2015; El-Brolosy et al., 2019; Ma et al., 2019). *lzk^la232^* mutants, however, have a premature stop codon, which could trigger nonsense-mediated RNA decay and thus genetic compensation, perhaps explaining why *lzk^la232^* mutants did not have a strong motor axon regeneration defect on their own. The fact that a few double mutant motor axons were able to regenerate may suggest compensatory contributions from other MAP3Ks in the MLK family or parallel pro-regenerative pathways. We saw a similar pattern for the suppression of RB neuron regenerative sprouting (sprouting was increased in *dlk^la231^* animals, but it was more pronounced in *dlk^la231^ lzk^la232^* double mutants), and only found motor dendrite overgrowth in double mutants. Together, these results suggest a partially redundant or compensatory relationship between these two kinases in axon regeneration.

DLK promotes axon regeneration in many different types of neurons and organisms, but in larval *Drosophila* sensory neurons it is dispensable for dendrite regeneration, even though it is required for axon regeneration in the same cells (Stone et al., 2014). This observation indicates that axon and dendrite regeneration are mechanistically distinct, and indeed other factors differentially affect these processes (Nye et al., 2020). Vertebrate sensory neurons, while similar in many respects to *Drosophila* counterparts, have sensory axons rather than dendrites, as defined by physiological features and cytoskeletal organization (Shorey et al., 2021), so we initially hypothesized that DLK and LZK would be required for regeneration of both central and peripheral sensory axons. Surprisingly, however, central RB, peripheral RB, and peripheral DRG axons regenerated normally in *dlk^la231^ lzk^la232^* mutants. These findings suggest that another, yet unknown pathway is used by fish somatosensory neurons to detect axon damage and activate the axon regeneration program.

When an entire nerve is damaged, it is difficult to deconvolute axon regeneration from a severed axon stump, regenerative sprouting from spared branches, and collateral sprouting from neighboring neurons (Steward et al., 2003; Tuszynski and Steward, 2012). The balance of these types of growth, however, has important functional consequences, particularly for sensory neurons. For example, RB sensory axons in larval zebrafish tile to innervate discrete, minimally overlapping territories in the skin that provide spatial information necessary for appropriate behavioral responses to touch (Sagasti et al., 2005). The balance between true regeneration and collateral sprouting thus determines the sensory map of the periphery. Our single cell labeling method allowed us to directly address this issue, revealing that DLK, potentially with some contribution from LZK, are required to limit sprouting from uninjured axon branches, even though it did not affect regeneration from severed axon branches. This finding emphasizes that forming a growth cone in a damaged axon branch, which has experienced cytoskeletal disruption, calcium influx, and local mitochondrial dysfunction, is a distinct process from reactivating growth in a dormant, uninjured axon branch. Since it is critical for sensory neurons to restore their spatial sensory map in the periphery, limiting sprouting may be as functionally important as promoting new growth from an injured branch. Indeed, excessive neuronal sprouting induced by pathological conditions such as atopic dermatitis, or selective serotonin reuptake inhibitors (SSRI) treatment, is associated with oversensitivity and itch in the skin (Han and Dong, 2014; Tominaga and Takamori, 2014; Morita et al., 2015).

The cellular site in which DLK acts to limit sprouting remains unclear in our experiments, since expressing DLK cDNA only modestly reduced sprouting in *dlk^la231^* RB neurons. Achieving rescue experimentally may be difficult, since overexpressing DLK was toxic to neurons, suggesting that precise regulation of DLK levels is required for an optimal injury response. These findings also allow the intriguing possibility that DLK may act non cell-autonomously to limit sprouting. For example, DLK could function in the epidermal cells innervated by RB axons, since DLK regulates epidermal differentiation and integrity (Robitaille et al., 2005; Simard-Bisson et al., 2017), or in immune cells activated by the injury, since DLK and LZK regulate microglial and astrocyte responses to injury in the CNS (Chen et al., 2018; Wlaschin et al., 2018). Wherever DLK is functioning to limit sprouting, our findings reveal that regeneration from an injured growth cone and sprouting from uninjured axon branches are mechanistically distinct processes that differentially require DLK signaling.

Our discovery that zebrafish DLK and LZK are required to promote axon growth in one context (motor axon regeneration), and restrain it in others (growth of RB spared branches and trigeminal neurons) echoes the context-specific dual roles for DLK homologs in other neurons. For example, in addition to its differential effects on axon and dendrite regeneration in *Drosophila* sensory neurons (Stone et al., 2014), excess DLK promotes axon growth and inhibits dendrite growth in the same neurons (Wang et al., 2013). Moreover, although DLK is required for axon regeneration in *Drosophila* sensory neurons, it also promotes an opposing neuroprotective response that inhibits axon regeneration (Chen et al., 2016a). Thus, DLK has the potential to both activate and inhibit regeneration in these neurons, depending on the balance of its outputs. Similarly, in *C. elegans* sensory neurites, excess DLK resulting from loss of a negative regulator can inhibit axon regeneration (Yan et al., 2009) or promote developmental overgrowth (Zheng et al., 2020), depending on which regulator was reduced.

Our study adds to a growing list of examples of distinct DLK functions in different cell types and conditions (Jin and Zheng, 2019). These diverse outcomes could be explained by the permissiveness versus non-permissiveness of the environment, expression levels of these signaling proteins, subcellular localization, the mode of activation, or even the duration of the signal, all of which could lead to the assembly of different signaling complexes that activate different responses (Goodwani et al., 2020). Understanding these diverse molecular processes, categorizing potential outcomes, and determining the neuronal cell types in which they occur in vivo, are prerequisites to considering *dlk* and *lzk* as targets in the treatment of axonal neuropathies and traumatic injuries.

## Materials and methods

### Zebrafish

Zebrafish (*Danio rerio*) were raised on a 14h/10h light/dark cycle, and a water temperature of 28.5C. Embryos were incubated at 28.5C in E3 buffer (0.3g/L Instant Ocean salt, 0.1% methylene blue). For imaging purposes, pigment formation was blocked by treating embryos with phenylthiourea (1X PTU, 0.2mM) at 22-24hpf. Embryos were then manually dechorionated using forceps. All mutant and transgenic lines were created using AB wildtype fish (ZFIN: ZDB-GENO-960809-7). Experimental procedures were approved by the Chancellor’s Animal Research Care Committee at UCLA and the Pennsylvania State Institutional Animal Care and Use Committee.

### CRISPR/Cas9 mutagenesis

Guide RNAs were engineered using the “Short oligo method to generate gRNAs”, as previously described (Talbot and Amacher, 2014). To mutagenize *dlk* and *lzk*, we created a DNA template for making gRNAs, containing the T7 RNA polymerase promoter, the gene targeting sequence, and the gRNA scaffold sequence. This template was amplified by PCR and its product was used to synthesize gRNAs using a T7 RNA polymerase kit. The gRNAs were then purified. Fish were injected with a mix containing 1uL of Cas9 mRNA (1500ng/ul), 1.5uL of gRNA1 (100ng/uL), and 1.5uL of gRNA2 (100ng/uL). Embryos at the 1-cell stage were injected with 5nL of the mix. At 48hpf, PCR and restriction digests were used to test guide efficiency. To identify fish carrying mutations in their germline, injected embryos were raised to adulthood, crossed to wildtype fish, and their progeny were screened by PCR. at 48hpf, mutant PCR products flanked by the common M13 primers, were sent for sequencing.

### PCR genotyping

PCR was usually conducted with Taq polymerase for 40 amplification cycles. The denaturation cycle was 94°C for 30s. Annealing was for 30s at 53°C for *lzk* wildtype and *lzk^la232^* bands, 59°C for the dlk wildtype band, and 63°C for *dlk^la231^* band. Elongation was at 72°C; this step was 20s for *lzk* wildtype and *lzk^la232^* bands, 1 minute for the *dlk* wildtype band, and 15s for the *dlk^la231^* band.

Alternatively, PCR was occasionally conducted with Phusion polymerase for 40 amplification cycles. The denaturation cycle was 98°C for 30s. Annealing was at 64°C for lzk wildtype and *lzk^la232^* bands, 71°C for the dl*k* wildtype band, and 75°C for *dlk^la231^* band. Elongation at 72°C for 11s for *lzk* wildtype and *lzk^la232^* bands, for 30s for the *dlk* wildtype band, and 10s for *dlk^la231^* band.

### Transgene cloning

#### HB9(3X)-E1B-DLK-T2A-GFP

E1B-DLK-T2A was constructed by individually inserting E1B and DLK into the MCS region of a PME: MCS-T2A vector.

*Step 1:* E1B template

5’ TCTAGAGGGTATATAATGGATCCCATCGCGTCTCAGCCTCA 3’

5’ GAATTCGTGTGGAGGAGCTCAAAGTGAGGCTGAGACGCGATG 3’

An E1B template was created by PCR amplification of overlapping oligomers.

*Step 2:* att site-MCS-T2A-att site

5’ GGGGACAAGTTTGTACAAAAAAGCAGGCT<u>ACCGTCAGATCCGCTAG</u> 3’

5’ GGGGACCACTTTGTACAAGAAAGCTGGGTA<u>TGGGCCAGGATTCTC</u> 3’

MCS-T2A flanked by att sites was PCR amplified and inserted into a PME vector using a Gateway BP reaction (Kwan et al., 2007).

*Step 3:* EcorI-E1B-EcorI was inserted into the MCS region of PME: MCS-T2A vector using restriction digest and ligation resulting in aPME: MCS(E1B)-T2A vector.

5’ TAAGCAGAATTCTCTAGAGGGTATATAATGGATCCCA 3’

5’ TGCTTAGAATTCGAATTCGTGTGGAGGAGCT 3’

*Step 4:* SaLI-DLK(no stop codon)-SacII was inserted into the PME: MCS(E1B)-T2A vector using restriction digest and ligation resulting in a PME: MCS(E1B-DLK)-T2A vector.

5’ TAAGCAGTCGACATGGCTTGTGTCCATGAGCAG 3’

5’ TGCTTACCGCGGGTTTTGTGGACCCTGGCCC 3’

PME: MCS(E1B-DLK)-T2A, a P5E:HB9(3X), and P3E:GFP were incorporated into a Gateway destination vector via LR reaction resulting in HB9(3X)-E1B-DLK-T2A-GFP.

#### HB9(3X)-E1B-LZK-T2A-GFP

E1B-LZK-T2A was constructed using overlap PCR to assemble E1B-LZK-T2A framed by att sites:

*Primer set 1:* att site-E1B-part of *lzk,* use E1B sequence as a template

5’ GGGGACAAGTTTGTACAAAAAAGCAGGCTTC<u>TCTAGAGGGTATATAATGGATCCC</u> 3’

5’ TGGTGCTGTGCGTGTGCAT<u>GAATTCGTGTGGAGGAGC</u> 3’

*Primer set 2:* part of E1B-*lzk* no stop codon-part of T2A, use *lzk* cDNA as template

5’ GCTCCTCCACACGAATTC<u>ATGCACACGCACAGCACCA</u> 3’

5’ CCTCTGCCCTCTCCACTTCC<u>CCAGGATGACGGAGCGCC</u> 3’

*Primer set 3:* end of *lzk* no stop codon-T2A-att site, use T2A sequence as a template

5’ GGCGCTCCGTCATCCTGG<u>GGAAGTGGAGAGGGCAGAGG</u> 3’

5’ GGGGACCACTTTGTACAAGAAAGCTGGGTC<u>TGGGCCAGGATTCTCCTCGA</u> 3’

All three fragments were amplified independently. Fragment 1 was added to fragment 2 by overlapping PCR. The resulting fragment was added to fragment 3. The complete sequence was then inserted into Gateway’s PME donor plasmid via a BP reaction. The resulting PME: E1B-LZK-T2A, a P5E:HB9(3X), and P3E:GFP were incorporated into a Gateway destination vector via LR reaction resulting in HB9(3X)-E1B-LZK-T2A-GFP.

#### Crest3: DLK-T2A-GFP

DLK-T2A was constructed using overlap PCR to assemble DLK-T2A flanked by att sites for Gateway recombination :

*Primer set 1:* att site-*dlk* no stop-part of T2A, use *dlk* cDNA as template

5’ GGGGACAAGTTTGTACAAAAAAGCAGGCTTC<u>ATGGCTTGTGTCCATGAGCAG</u> 3’

5’ CCTCTGCCCTCTCCACTTCC<u>GTTTTGTGGACCCTGGCCC</u> 3’

*Primer set 2:* end of *dlk* no stop-T2A-att site, use T2A sequence as template

5’ GGGCCAGGGTCCACAAAAC<u>GGAAGTGGAGAGGGCAGAGG</u> 3’

5’ GGGGACCACTTTGTACAAGAAAGCTGGGTC<u>TGGGCCAGGATTCTCCTCGA</u> 3’

Both fragments were amplified independently. Fragment 1 was added to fragment 2 by overlapping PCR. The resulting PME: DLK (no stop codon)-T2A, a P5E: Crest3, and a P3E:GFP were incorporated into a Gateway destination vector via LR reaction resulting in Crest3: DLK-T2A-GFP.

## Building the phylogenetic tree

Complete DLK and LZK protein sequences from several organisms were downloaded from the NCBI database. The sequences were aligned using the MUSCLE alignment algorithm on the EMBL-EBI website. Phylogenetic analyses were performed with the RAxML software using a maximum likelihood method, the JTT substitution matrix, and empirical frequencies (Stamatakis, 2014). The RAxML software was accessed via the CIPRES Science Gateway (Miller et al., 2010). The Interactive Tree of Life website (Letunic and Bork, 2007) was used to visualize the evolutionary tree.

### Larval body measurements

The body lengths of larvae of each genotype were measured at 48 hpf, 72 hpf and 5 dpf using a Zeiss Discovery V8 Stereomicroscope at a 2X magnification. Bodies were measured lengthwise from head to tail, and tail widths measured ventral to dorsal at the end of the spinal cord. Larvae from different clutches (n>8) and parents of different ages (3-24 months old) were mixed for this analysis.

### Mounting larvae for live imaging

Larva were anesthetized with 0.2 mg/mL MS-222 in embryo media (0.08%) before mounting. Each larva was embedded in 1% agarose and placed on a cover slip. Upon solidification of the agarose (∼15 minutes), a plastic ring was sealed onto the cover slip with vacuum grease. The resulting chamber was then filled with tricaine-containing embryo media and sealed with a glass slide using vacuum grease (O’Brien et al., 2009b).

### Larval microscopy

Live confocal images were collected on an LSM 800 using a 20X air objective (Plan-APOCHROMAT, NA= 0.8). Images were acquired with Zen Blue software from Zeiss.

Laser axotomies were performed using a LSM 880 equipped with a 2-photon laser (Chameleon Ultra II, Coherent), as previously described (O’Brien et al., 2009b). Zen Black 2.1 SP3 software was used to visualize axons and perform axotomies. Neurons were visualized with a 561nm or 488nm laser excitation, before switching to the Chameleon (813nm) laser for axon severing.

To cut axons of RB neurons in the tail, we initially used 5% laser power. If axons were not cut, the laser power was increased in 0.5% increments until cutting was successful. Since MNs are deeper in the animal, we first attempted axotomy with 6.5% laser power.

### Adult experiments

Cutting DRG nerves in juvenile fish: 4-5 week-old fish of the genotype P2rx3a:LexA,4xLexOP:EB3-GFP (not shown in the figure), P2rx3a:LexA,4xLexAop:mCherry^la207^, Roy^a9^/Roy^a9^, mifta^w2^/mifta^w2^, containing either wt or homozygous *dlk*^la231^ and *lzk^la232^* were anesthetized in a VWR polystyrene petri dish, filled halfway with 0.16% tricaine in 0.6 g/L Instant Ocean salt solution, and immobilized by applying agarose to their midsection only, leaving both the head’s respiratory apparatus, and tail free. Fish were then imaged on a Leica SP8 microscope equipped with an InSight X3 unit from Spectra-Physics. A 25X (NA= 1) water immersion objective with a working distance of 2.6mm was used to image the posterior spinal cord of the fish, and an ROI was chosen to restrict the cut site to the width of the nerve and positioned ∼100 μm from the posterior-most DRG, along the posterior projecting nerve. The tunable laser was set to 900nm, and both the tunable and fixed wavelength 1045nm lasers were set to 100% on the slowest speed setting and scanned for ∼1 second. The 2-photon overview showing the nerve stumps after cut was performed on the aforementioned SP8, all other images for this experiment were obtained with a Zeiss LSM 800 Axio Observer Z.1 with a 20X air objective (NA=0.8).

Cutting scale nerves: All fish were between 8 and 11 months old. The single *dlk^la231^* and *lzk^la232^* mutants were transgenic for P2rx3a:LexA,4xLexAop:mCherry^la207^, and did not possess mutant roy or mifta alleles. Wildtype and double mutant fish were in a roy^a9^/roy^a9^, mifta^w2^/mifta^w2^ background and doubly transgenic for both P2rx3a:LexA,4xLexAop:EB3-GFP and P2rx3a:LexA,4xLexAop:mCherry^la207^. All images shown were collected on a Zeiss LSM 800 Axio Observer Z.1. For wildtype, a 25X a multi-immersion objective was used (NA= 0.8), and for the double mutants a 20X air objective (NA= 0.8) was used. Laser injury was performed using an Andor Micro-Point UV pulse laser.

### Image analysis and statistics

Confocal images, saved as .czi files, were opened in ImageJ/Fiji and measured using the Simple Neurite Tracer (SNT) feature in z-stack format. One neuron was imaged per embryo. Axons of MNs were traced from the cell body to their endings in muscles. RB peripheral arbors were traced starting at the first branch point in the skin. If RB peripheral axons bifurcated in the spinal cord, creating two separate peripheral arbors, both arbors were traced. Three experimenters contributed to tracing. To test for tracing reproducibility between experimenters, a subset of RB central axons (n=76) and peripheral arbors (n=10) were separately traced by two experimenters. In both cases, tracings by the two experimenters were highly similar (r=0.93 for the central axons; r=0.99 for the peripheral arbors).

For figures, maximum projections were created in ImageJ/Fiji, converted to grayscale and inverted. To visualize entire RB neurons, images of different parts of each neuron were stitched together in Adobe Photoshop.

Dot plots, or box-and-whisker plots overlaid with dot plots were produced in R to visualize the data. In dot plots, the black bar indicates the mean. In box-and-whisker plots, the boxes indicate the interquartile range: the top of the box is the 75th percentile, the midline is the median, and the bottom of the box is the 25th percentile. Statistical analyses were performed in R. A generic quantile-quantile test was used to determine the normality of sample populations. Unless otherwise specified, data distributions were non-parametric. Therefore, a Kruskal-Wallis, with a Bonferroni correction, was used to assess differences between experimental groups. A Wilcoxon paired test was used to identify groups with significant differences. * for a p-value < 0.05, ** for a p-value < 0.01, *** for a p-value < 0.005, and **** for a p-value < 0.001.

**Table 1:** Accession numbers

| **DLK** | Accession number | Common name |
| --- | --- | --- |
| *Homo_sapiens_DLK* | NP_006292.3 | Human |
| *Rattus_norvegicus_DLK* | NP_037187.1 | Rat |
| *Mus_musculus_DLK* | NP_001345773.1 | Mouse |
| *Gallus_gallus_DLK* | XP_025001292.1 | Chicken |
| *Danio_rerio_DLK* | NP_996977.1 | Zebrafish |
| Xenopus_laevis_DLK | NP_001094411.1 | Frog |
| *Caenorhabditis_elegans*_DLK-1 | sp\|O01700.4 | Worm |
| *Drosophila_megalogaster*_wallenda | NP_788541.1 | Fruit fly |
| **LZK** |  |  |
| *Homo_sapiens*_LZK | sp\|O43283.1 | Human |
| *Rattus_norvegicus*_LZK | NP_001014000.2 | Rat |
| *Mus_musculus*_LZK | NP_766409.2 | Mouse |
| *Gallus_gallus*_LZK | XP_025009526.1 | Chicken |
| *Danio_rerio*_LZK | XP_017213349.1 | Zebrafish |
| Xenopus_laevis_LZK | ABK15544.1 | Frog |

**Table 2:**
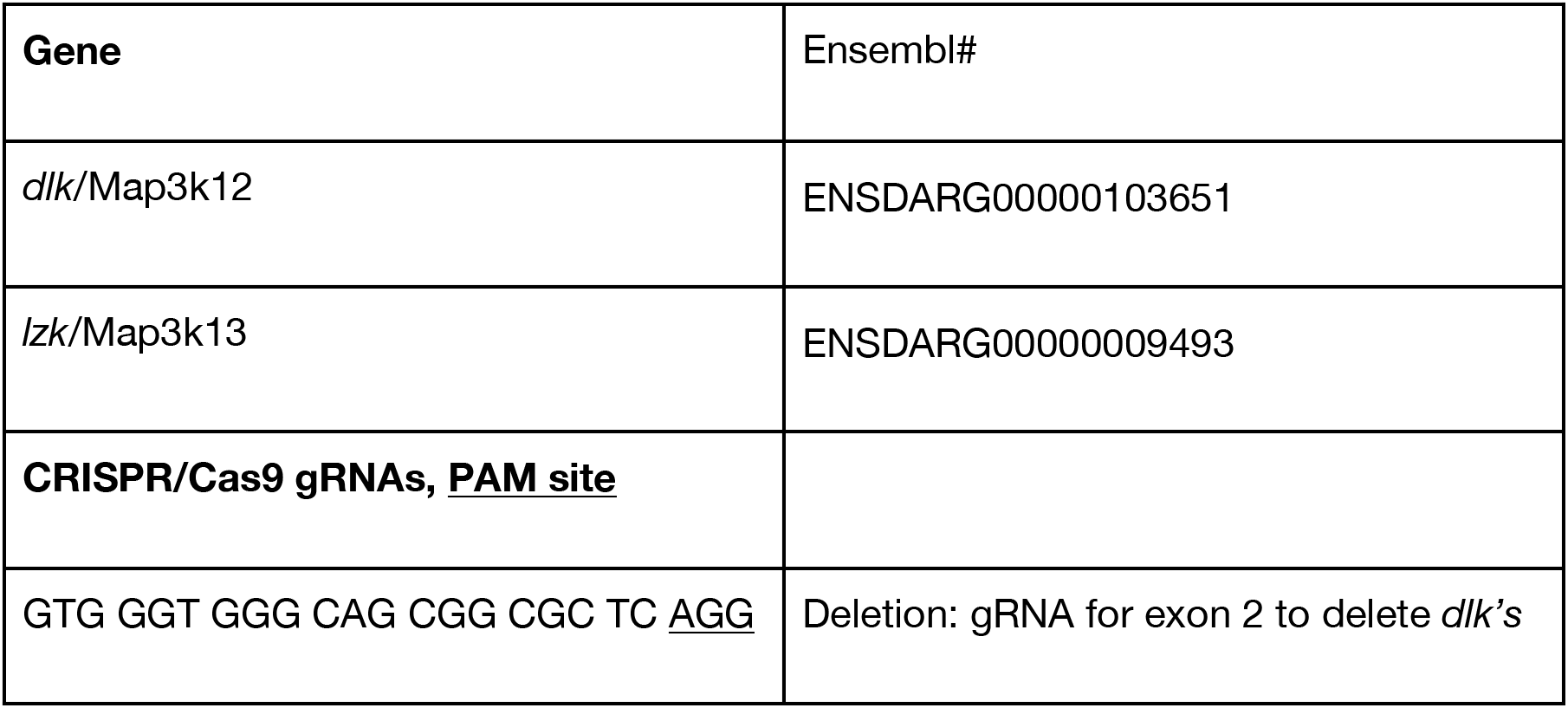
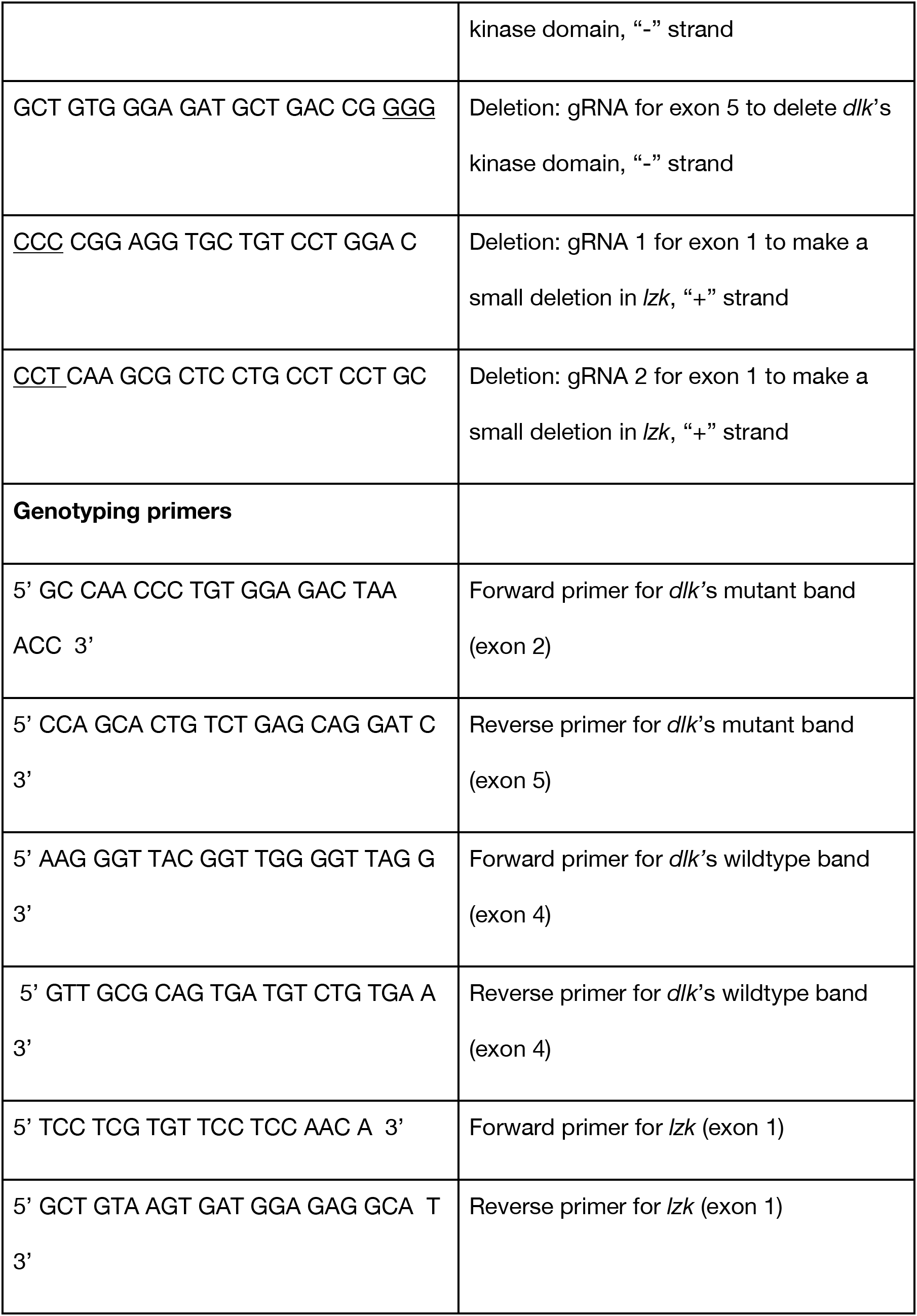
Gene IDs, Cas9 gRNAs, genotyping primers

**Table 3:**
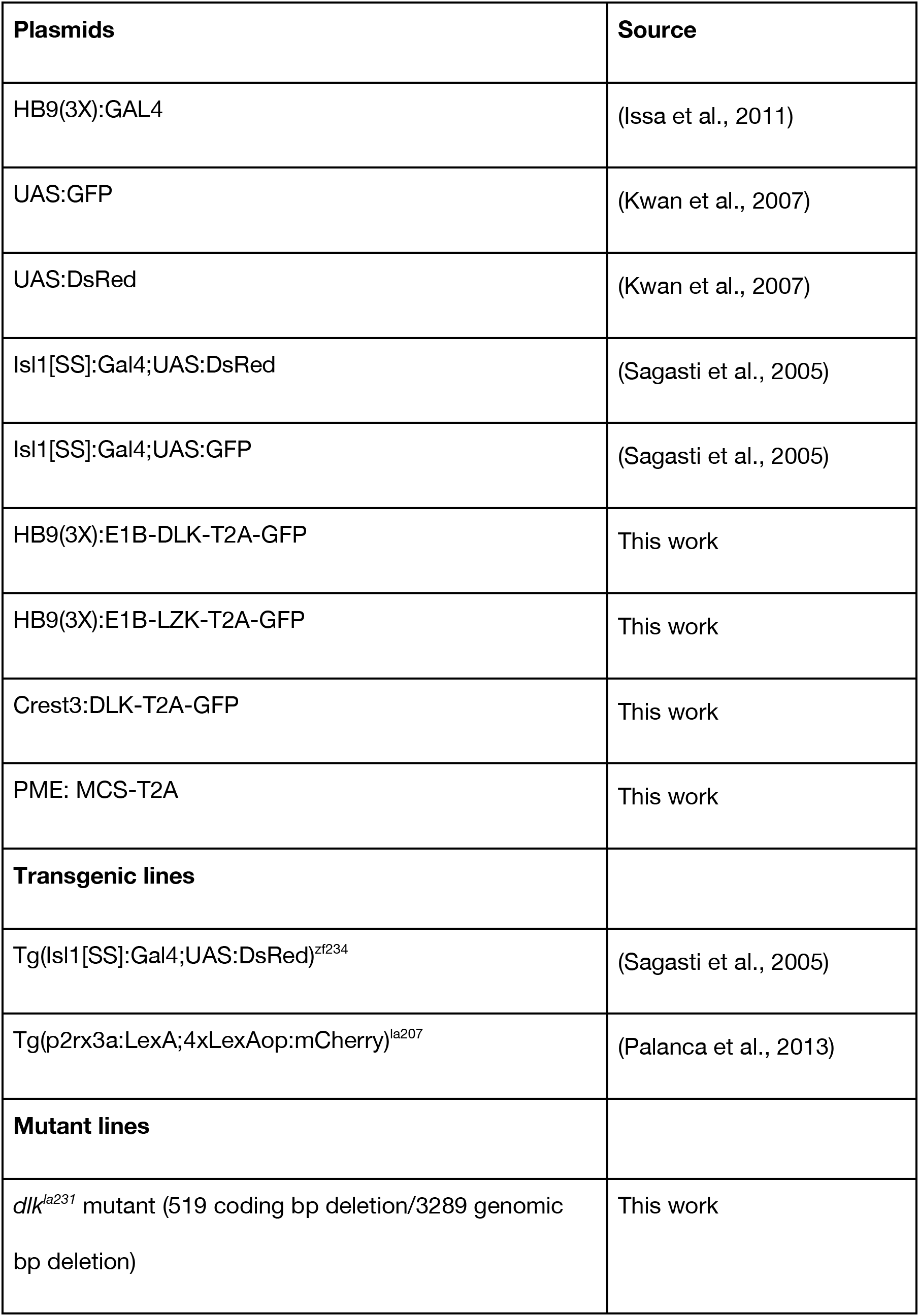
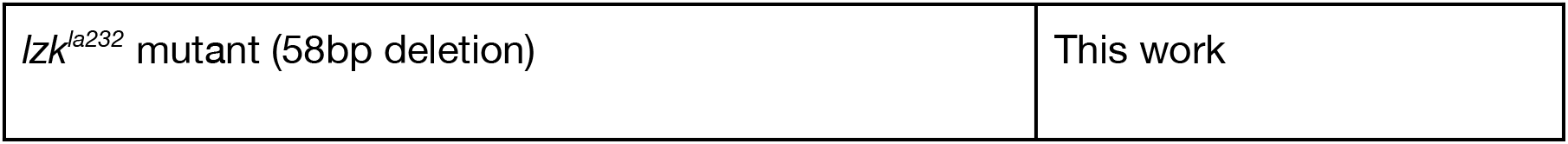
Plasmids used in injections, transgenic lines

| **Plasmids** | **Source** |
| --- | --- |
| HB9(3X):GAL4 | (Issa et al., 2011) |
| UAS:GFP | (Kwan et al., 2007) |
| UAS:DsRed | (Kwan et al., 2007) |
| Isl1[SS]:Gal4;UAS:DsRed | (Sagasti et al., 2005) |
| Isl1[SS]:Gal4;UAS:GFP | (Sagasti et al., 2005) |
| HB9(3X):E1B-DLK-T2A-GFP | This work |
| HB9(3X):E1B-LZK-T2A-GFP | This work |
| Crest3:DLK-T2A-GFP | This work |
| PME: MCS-T2A | This work |
| **Transgenic lines** |  |
| Tg(Isl1[SS]:Gal4;UAS:DsRed)<sup>zf234</sup> | (Sagasti et al., 2005) |
| Tg(p2rx3a:LexA;4xLexAop:mCherry)<sup>la207</sup> | (Palanca et al., 2013) |
| **Mutant lines** |  |
| *dlk*<sup>la231</sup> mutant (519 coding bp deletion/3289 genomic bp deletion) | This work |
| *lzk*<sup>la232</sup> mutant (58bp deletion) | This work |

**Table 4:** Reagents, resources

| Reagents | Source |  |
| --- | --- | --- |
| HiScribe T7 High Yield RNA Synthesis Kit | New England Biolabs | E2040S |
| RNeasy Mini Kit | Qiagen | 74104 |
| Alt-R S.p. Cas9 Nuclease 3NLS | IDT | 1074182 |
| PfuUltra II Fusion HotStart DNA Polymerase | Agilent | 600670 |
| Phusion High-Fidelity DNA Polymerase | New England Biolabs | M0520S |
| *Taq* DNA Polymerase with Standard *Taq* Buffer | New England Biolabs | M0273X |
| Zen 2.1(Blue edition) | Carl Zeiss Microscopy | http://www.zeiss.com |
| Fiji/ImageJ | (Schindelin et al., 2012) | https://fiji.sc/ |
| R | R Foundation for Statistical Computing | https://www.r-project.org/ |

## Acknowledgments

We thank members of the Sagasti lab for comments on the manuscript, Son Giang and Yuan Dong for excellent fish care, and Adam Langenbacher for help with data presentation. KPA was supported by a UCLA Cota-Robles fellowship, a NSF Bridge to the Doctorate fellowship, and NIH F31 predoctoral fellowship NS106742. This study was supported by NIH grants R21NS090027 (MMR and AS) and R01AR064582 (AS).

---

**Supplemental Figure 1.**
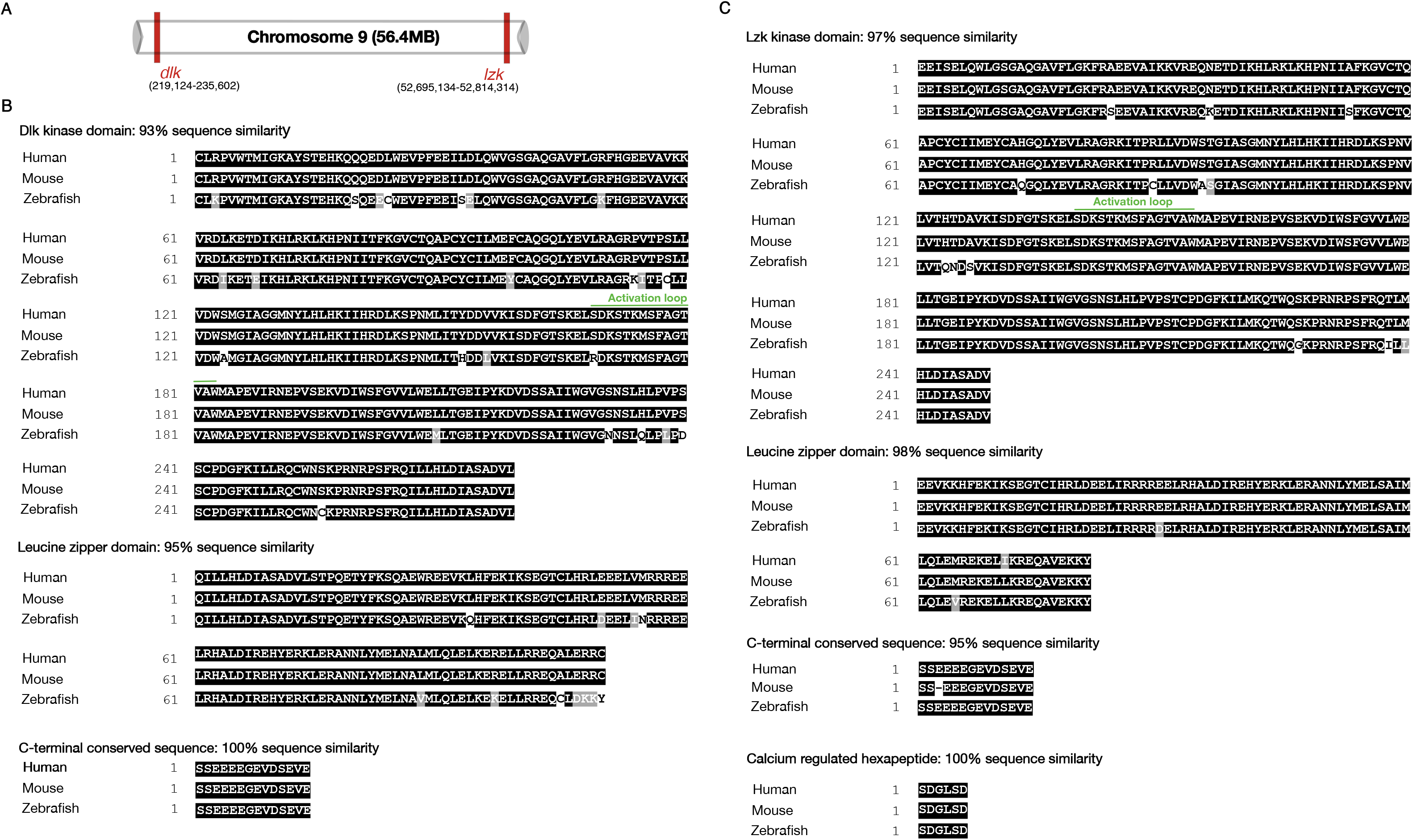
Zebrafish *dlk* and *lzk* genes and alignment of key domains. A) Depiction of *dlk* and *lzk* locations on chromosome 9. B) Alignment of human, mouse, and zebrafish DLK kinase, leucine zipper, and C-terminal conserved protein sequences. B) Alignment of human, mouse, and zebrafish LZK kinase, leucine zipper, C-terminal conserved sequence, and hexapeptide repeat protein sequences. Activation loops of the kinase domains are highlighted.

**Supplemental Figure 2.**
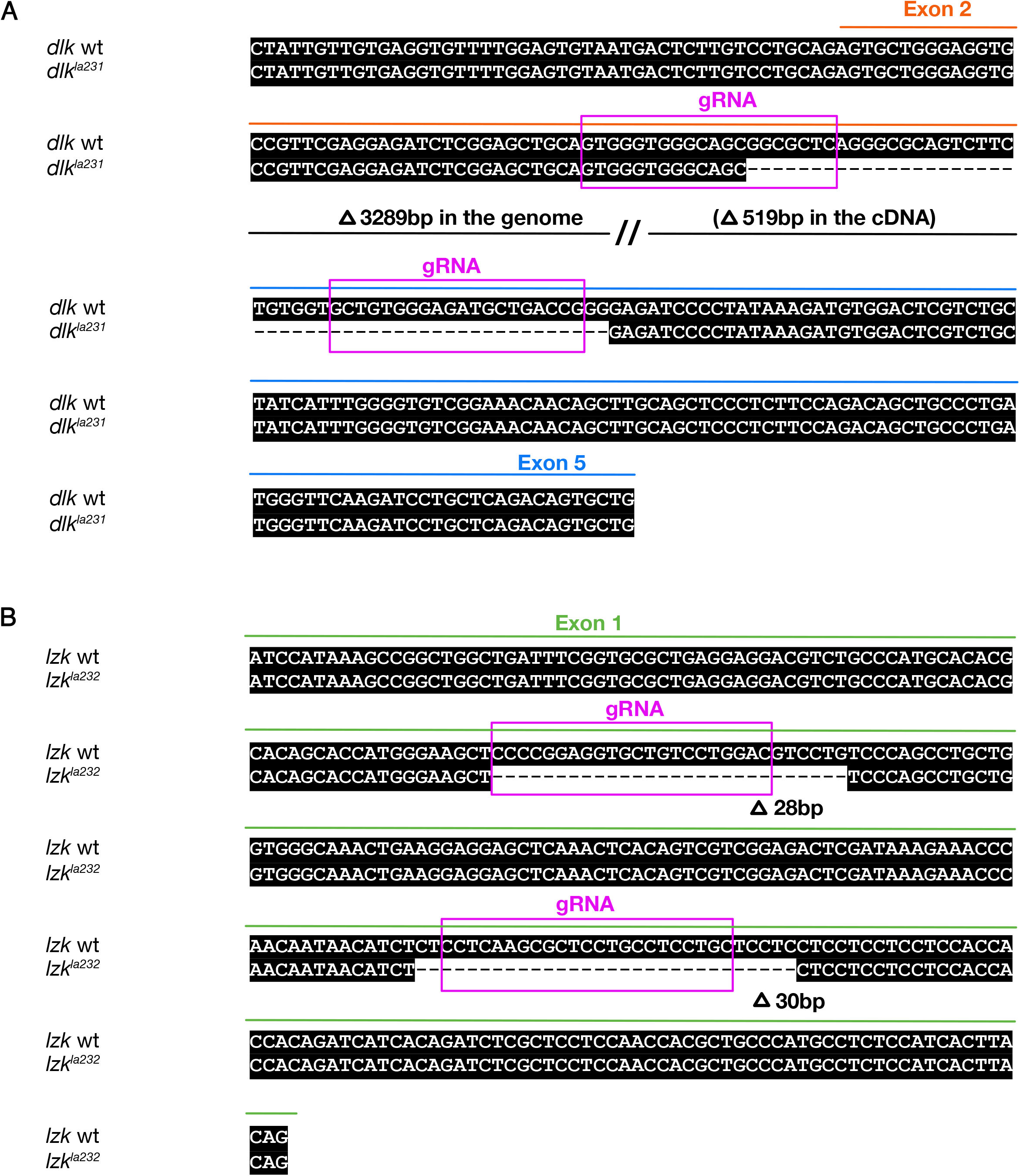
Diagram of *dlk^la231^* and *lzk^la232^* mutant sequences. A) Alignment of wt and *dlk^la231^* mutant DNA sequences showing deletion location. Exons and guide RNA sequences are indicated. B) Alignment of wt and *lzk^la232^* mutant DNA sequences showing deletion location. Exons and guide RNA sequences are indicated.

**Supplemental Figure 3.**
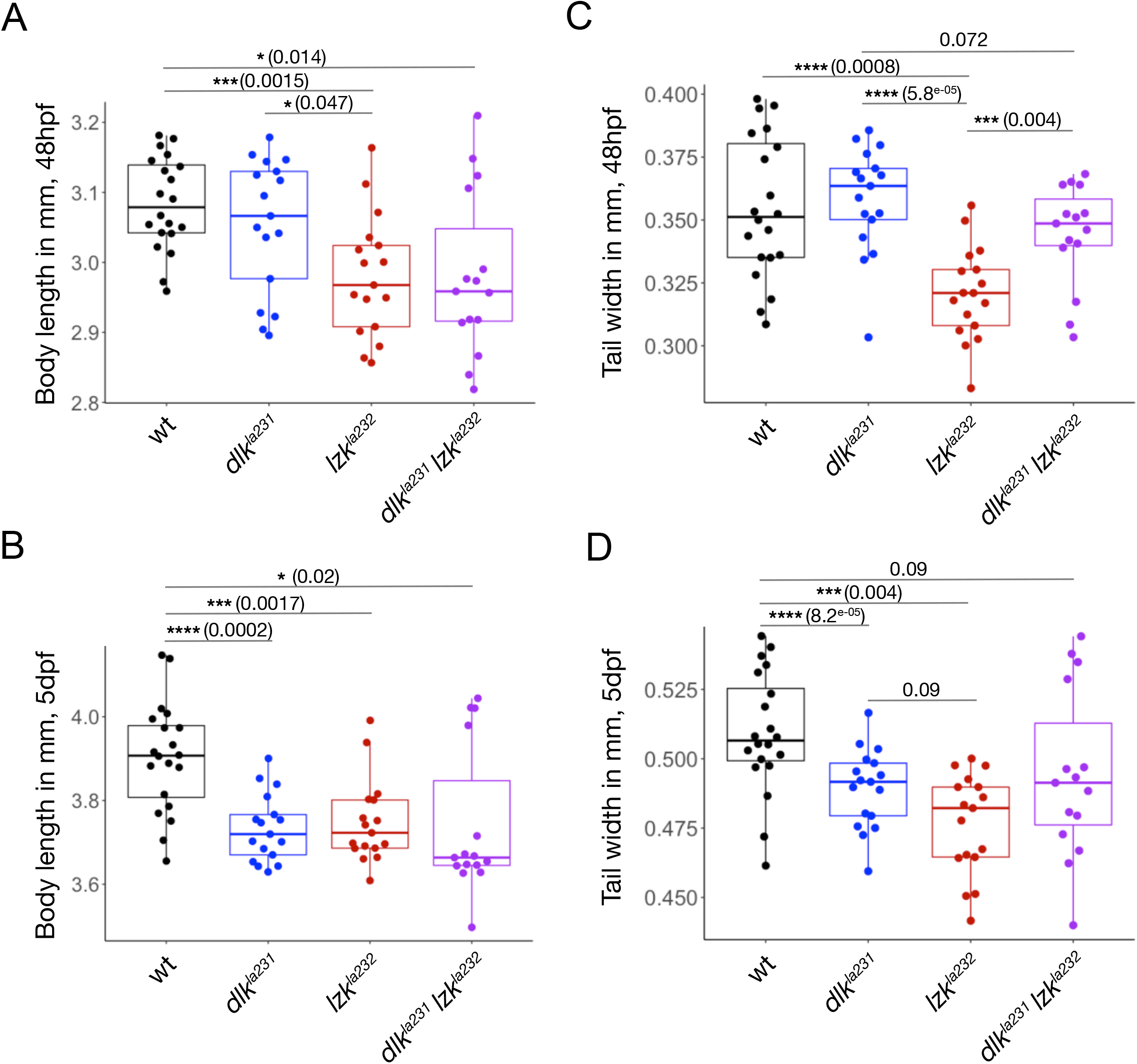
Quantification of larval body measurements at additional time-points. Overlaid box and dot plots showing body length and tail width at 48hpf and 5dpf in the indicated genotypes. See Methods for details of statistical analyses.

**Supplemental Figure 4.**
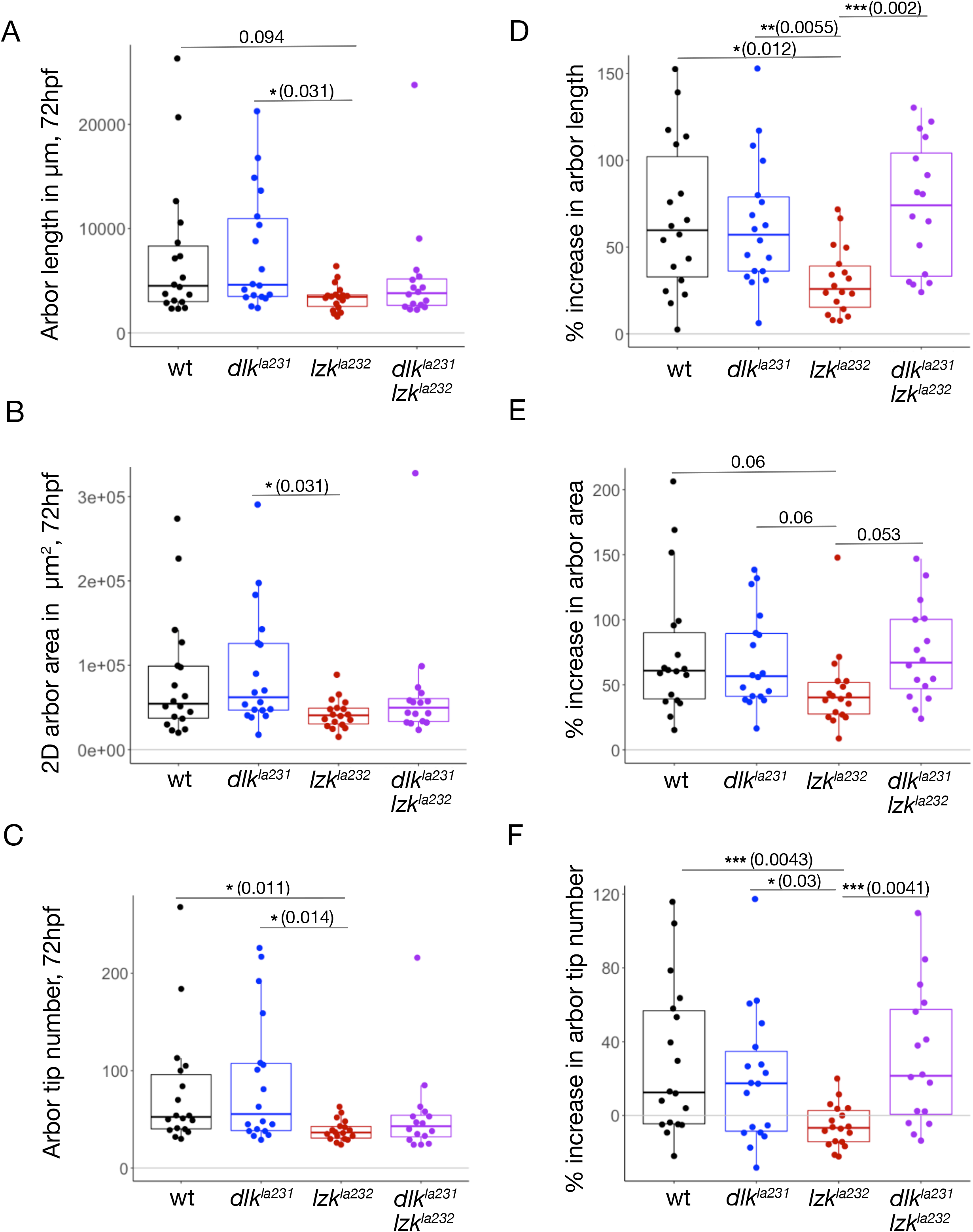
Additional quantification of peripheral axon morphology in uninjured RB neurons. A-C) Overlaid box and dot plots showing arbor length (A), 2D arbor area (B), and tip number (C) at 72hpf. D-F) Overlaid box and dot plots showing percent increase in arbor length (D), 2D area (E), and tip number (F) from 48 to 72hpf. See Methods for details of statistical analyses.

**Supplemental Figure 5.**
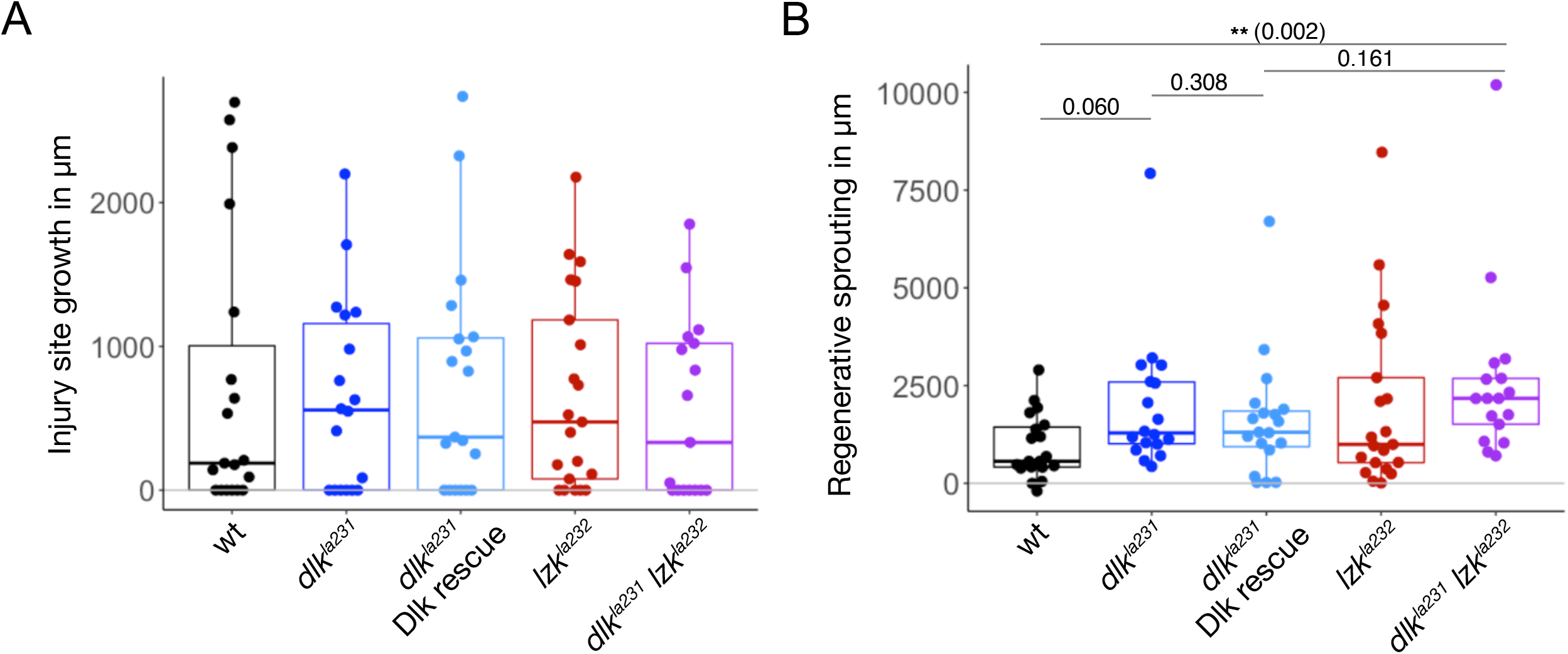
Additional quantification of axon growth in partial RB axotomies. A-B) Overlaid box and dot plots showing total length of post-axotomy growth at the injury site (A) and at the spared branch (regenerative sprouting) (B). See Methods for details of statistical analyses.

## Notes

### Competing Interest Statement

The authors have declared no competing interest.

